# Return of the Tbx5; lineage-tracing reveals ventricular cardiomyocyte-like precursors in the injured adult mammalian heart

**DOI:** 10.1101/2022.01.28.473428

**Authors:** Panagiota Siatra, Giannis Vatsellas, Athanasia Chatzianastasiou, Evangelos Balafas, Theodora Manolakou, Andreas Papapetropoulos, Anna Agapaki, Eleni-Taxiarchia Mouchtouri, Artemis G. Korovesi, Manolis Mavroidis, Dimitrios Thanos, Dimitris Beis, Ioannis Kokkinopoulos

## Abstract

The single curative measure for heart failure patients is total heart transplantation, which is limited due to a shortage of donors, the need for immunosuppression and economic costs. Therefore, there is an urgent unmet need for identifying cell populations capable of cardiac regeneration that we will be able to trace and monitor. Injury to the adult mammalian cardiac muscle, often leads to a heart attack through the irreversible loss of a large number of cardiomyocytes, due to an idle regenerative capability. Recent reports in zebrafish indicate that Tbx5a is a vital transcription factor for cardiomyocyte regeneration. Preclinical data underscore the cardioprotective role of Tbx5 upon heart failure. Data from our earlier murine developmental studies have pinpointed a prominent unipotent Tbx5-expressing embryonic cardiac precursor cell population able to form cardiomyocytes, *in vivo*, *in vitro* and *ex vivo*.

Using a developmental approach to an adult heart injury model and by employing a lineage-tracing mouse model as well as the use of single-cell RNA-seq technology, we identify a Tbx5-expressing ventricular cardiomyocyte-like precursor population, in the injured adult mammalian heart. The transcriptional profile of that precursor cell population is closer to that of neonatal than embryonic cardiomyocyte precursors. Tbx5, a cardinal cardiac development transcription factor, lies in the centre of a ventricular adult precursor cell population, which seems to be affected by neurohormonal spatiotemporal cues.

The identification of a Tbx5-specific cardiomyocyte precursor-like cell population, which is capable of dedifferentiating and potentially deploying a cardiomyocyte regenerative program, provides a clear target cell population for translationally-relevant heart interventional studies.

## INTRODUCTION

According to the World Health Organisation, heart failure is the major cause of death in industrialised countries with an estimated 17.3 million deaths per year due to cardiovascular disease, representing 30% of all global deaths^1^. Human heart regeneration is one of the most critical unmet clinical needs at a global level. Congenital heart defects (CHDs) are usually apparent at birth, characterised by structural abnormalities, such as atrial or ventricular septation defects, electrical conduction abnormalities or cardiomyopathies. One of the primary causes of cardiomyopathies is the loss and/or damage of heart muscle cells, termed cardiomyocytes (CM). In order to replenish the lost/damaged cells, an appropriate source is needed as a cell-replacement therapeutic approach. An attractive candidate is cardiac precursor cells (CPCs), which could be driven to give rise to mature CMs.

During development, the CM lineage is highly specialised, comprising cardiac progenitors allocated in a discrete and temporal order^2^. At embryonic day (E) 7.5 in mice^3,4^, the heart tube is the initial structure that eventually gives rise to the heart proper. It is populated by two distinct sets of cardiac progenitors derived from two anatomical regions; the first heart field (FHF also known as the cardiac crescent) which will give rise to the left ventricle (LV) and parts of the atria, and the second heart field (SHF) that contributes towards the right ventricle, outflow tract and the remaining parts of the atria, including the septum^3–6^. These fields are genetically distinguishable, at E7.5, by expression of specific transcription factors (TF)^3,6,7^. After birth, most CMs are acytokinetic and have undergone terminal differentiation. However, recent studies have shown that the adult heart exhibits a capacity, albeit limited, to generate new CMs^8^. Carbon-14 birth-dating studies have suggested that around 40% of CMs are replaced over an entire life span, while IdU-labelling raises this percentage to 100%, in humans^9^, with a 1% per annum of CM turnover in the mammalian heart^10–13^. Tbx5, the T-box TF haploinsufficient in Holt-Oram syndrome, is one of the cardinal TFs essential for cardiac development and adult CM formation both *in vivo* and *in vitro*^14–18^.

Cardiomyocyte renewal in mammals could potentially be enabled *via* CM dedifferentiation and subsequent proliferation, as in the case of urodele amphibians and zebrafish^19^ Recent experiments performed in the adult zebrafish, reported that re-expression of *tbx5a* was essential for complete heart regeneration, upon ventricular ablation^20–22^. Thus, the Tbx5 transcriptional network is crucial not only for initiating early cardiac specification but to also prime the cardiac regenerative program, at least in adult lower vertebrates (^14^ and references within). While the priming of a resident CM proliferation program is now a leading therapeutic goal, the importance of Tbx5 in adult and postnatal mammalian heart ventricle regeneration has not been examined.

By employing a BAC *Tbx5^CreERT2/CreERT2^* transgene injury heart model^23^, we report the presence of cardiac cells that have over-activated Tbx5 following myocardial injury, with markers shown to be expressed in early cardiovascular precursors. Tbx5 transcription is controlled by a positive feedback loop in early murine heart development^23^. Therefore, Tbx5 overexpression upon injury could be one of the early attempts for priming regulatory networks important for CM dedifferentiation, division and/or differentiation in mice and humans^16–18^.

## METHODS

### Animals

The BAC transgene *Tbx5^CreERT2^* was constructed from the BAC clone RP23-267B15^24^ by replacement of exon 2 of Tbx5 with a CreERT2 cassette at the first methionine of the open reading frame in EL250 cells as described previously^23,25^. The *BAC-Tbx5^CreERT2^* transgenes were crossed with *Rosa26R^eYFP/eYFP^* transgenic mice (B6.129X1-Gt(ROSA)26Sortm1(EYFP)Cos/J) from Jackson laboratories stock# 006148, in order to produce *Tbx5^CreERT2^/Rosa26R^eYFP/eYFP^* mice employed in this study. The *Tbx5*^*CreERT2*/+^/*Rosa26R*^*eYFP*/+^/*Rosa26R*^*iDTR*/+^ was created using the tamoxifen-induced *Rosa26R^iDTR/iDTR^* transgene^26^ (Jackson laboratories stock# 007900) provided by the Klinakis lab (BRFAA).

All animal work has been approved by the BRFAA ethics committee and the Attica Veterinary Department (Animal Licence; 60876/23-1-20). All animals used were 2-3 months of age upon the time of LAD non-permanent ligation or isoproterenol administration experiments following relevant inclusion/exclusion guideline criteria^27^.

### Genotyping and PCR conditions

Genomic DNA extraction was performed from mouse tails with alkaline lysis (25mM NaOH, 0.2mM EDTA, pH=12 for 1.5h at 95°C) and pH was neutralized with Tris-HCL (pH=5) for 10 min RT. PCR conditions were as follow: initial denaturing step 3 min at 95°C, 35 cycles (15 s at 95°C, 15 s at 60°C and 58°C for Cre and eYFP primer pair respectively, 30 s at 72°C) and final extension 5 min at 72°C using KAPA Taq DNA polymerase (KAPA BIOSYSTEMS). PCR products were visualized on 2% agarose gels containing SYBR Safe DNA gel stain (Invitrogen). Primers used are;

CreRT2: F:5′-AGTTGCTTCAAAAATCCCTTCCAGGGCCCG-3′
R: 5′-AGCAATGCTGTTTCACTGGTTATGCGGCGG-3′
ROSA26 eYFP: F: 5′-GCGAAGAGTTTGTCCTCAACC-3′ R: 5′-AAAGTCGCTCTGAGTTGTTAT-3′
ROSA26 WT: 5′-GGAGCGGGAGAAATGGATATG-3′ R: 5′-AAAGTCGCTCTGAGTTGTTAT-3′

### Myocardial Infraction Models

Ischemia/reperfusion (I/R) injury; MI was induced in 2-3 month-old mice by a 10-minute transient ligation of the left anterior descending artery (LAD) as described previously^28^ with some modifications. Hearts were obtained 5 days after I/R (N=8).

Isoproterenol-induced cardiac infraction; Adult two-three month-old *Tbx5^CreERT2^/Rosa26R^eYFP/eYFP^*, *Tbx5*^*CreERT2*/+^/*Rosa26R*^*eYFP*/+^/*Rosa26R*^*iDTR*/+^ and control littermates were injected with isoproterenol (ISO, 20 mg/kg per day intraperitoneally, Sigma-Aldrich; St. Louis, I6504) once daily for two consecutive days^29–32^. Hearts were obtained and examined 2, 4, 6, 7 and 30 days after the last ISO injection (N=56).

Tamoxifen diluted in peanut oil (Sigma-Aldrich; St. Louis, P2144) was administrated on days 1 and 2 on all animals, at a final concentration of 0.8mg/10g of body weight, by oral gavage (N=84).

### Single cell RNA-Seq library preparation and Deep Sequencing

The Fluidigm C1 system was used to prepare single cells for RNA-Seq. RNA-Seq-IFCs were selected to capture all major cell populations from all cell size ranges observed using IFCs which capture cells of different sizes: 5-10uM (embryonic), 10-17uM (embryonic, neonatal), 17-25uM (>3 weeks of age). No batch effects were observed between chips of the same size. Onboard cell lysis, reverse transcription and cDNA synthesis were performed using the SMART-Seq v4 Ultra Low RNA Kit for the Fluidigm C1 System (Takara) reagents, following the manufacturer’s protocol. The resulting cDNAs from individual cells were used for the construction of NGS libraries with the Nextera XT DNA sample preparation kit (Illumina). Libraries were pooled, quantified with qubit HS DNA spectrophotometer and quality control was performed with the Agilent Bioanalyzer HS DNA kit. Approximately 1 Million 2×150bp Paired End Reads were generated for each single-cell RNA-Seq library in Illumina NovaSeq system following the manufacturer’s standard protocol. Count data were normalised to counts per million and transformed to Log2(CPM+1). Single-cell libraries with >500,000 reads and <5% in mitochondrial genes were used for further analysis.

### Single-Cell cDNA Expression Profiling

Embryonic murine heart cells were used from our previous studies and other research groups ^23,33^ for the purpose of comparing our *in vitro* embryonic and adult cardiac cell CPC transcriptomes. Embryonic FACS-sorted CPC on days 7 and 9 in vitro differentiation and adult CPC were collected *via* FACS sorting and further analysed using the Fluidigm C1 machine and workflow according to the manufacturer’s protocol. We examined a total of 20 cells derived from embryonic heart between E9.5-E10.5, 76 cells derived from P5 CPC, 22 Pdgfra^+^ interstitial adult cardiac cells and 240 YFP^+^ cells from D7 injured ventricles.

Part of the sequence data have already been submitted to NCBI Gene Expression Omnibus (GEO, http://www.ncbi.nlm.nih.gov/geo) under the accession number GSE63796.

### NGS data analysis pipeline

FASTQ data were quality tested and aligned to the murine genome using the online www.useGalaxy.eu platform. BAM files were created using the gapped-read mapper RNA-STAR alignment software using the mm10 murine primary assembly^34^. RNA-STAR parameters are depicted in **Supplementary Table 1.** BAM files were further sorted using Samtools sort^35^. Aligned and sorted BAM files were further analysed on SeqMonk 1.48.0 software as shown previously^36^ (**Supplementary Figure 7 and Supplementary Data 1**). For Gene Ontology (GO) and KEGG downstream analysis, toppgene online tool^37^, as well as cytoscape and ClueGo^38^ software, and STRING^™^ online tool were employed.

### Cardiac differentiation of murine ES cells

Cardiac differentiation of mouse ES cells (both *BAC-Tbx5^Cre^/R26R^eYFP/eYFP^* and BAC-*Tbx5^Cre^/Rosa26R^eYFP/eYFP^/Tbx5^KO/KO^* cell lines) was induced via monolayer and embryoid body conditions as previously described^23,39^. In brief, undifferentiated colonies were passaged for cell counting and re-seeded onto 24-well laminin-coated plates (5μg/ml) (Biolaminin LN521, BioLamina) pre-incubated with laminin for 2h at 37°C under 5% CO2, at a cell density of 120,000 cells/ well. Cells were cultured in ESGRO Complete medium plus LIF (Millipore 2i Medium Kit) for a day further. For three-step differentiation, cells were incubated for 1 day in IMDM/Ham’s F12 (Invitrogen) supplemented with N2 and B27 supplements (Gibco), 10% (stock) bovine serum albumin (Sigma), 2mM L-glutamine (Gibco), penicillin-streptomycin (Gibco), 0.5 mM ascorbic acid (Sigma), and 0.45 mM monothioglycerol (MTG, Sigma). For mesodermal induction and patterning, cells were exposed for 2 days to human Activin A (8 ng/ml, R&D Systems) and human bone morphogenetic protein 4 (hBMP4, 1.5 ng/ml; R&D Systems) together with human vascular endothelial growth factor (hVEGF, 5 ng/ml; R&D Systems). Cardiac specification was induced by exposure of cells to StemPro-34 SF medium (Gibco) supplemented with 2 mM L-glutamine, 0.5 mM ascorbic acid, human VEGF (5 ng/ml), human basic fibroblast growth factor (bFGF, 10 ng/ml, R&D Systems), and human fibroblast growth factor 10 (FGF10, 50 ng/ml, R&D Systems). The medium was changed every other day. All media were prepared under a sterile hood (Class 2), filtered through a Millipore Stericup 0.22mm filtration system and stored at 4 °C. Upon medium exchange, cells were washed twice with phosphate buffered saline (PBS, Gibco). Medium was changed every other day, and cells were analysed on various designated days. EBs (3000 cells per 30 μl in each drop) were dissociated for medium changes. 4-hydroxytamoxifen (4OH-TAM) administration (500nM) began on day 4 of differentiation and was renewed along with medium changes every other day.

### Cardiac Single-Cell Suspension Preparation

Adult hearts (and parts of) were collected and harvested on days 2, 4 and 7 after MI and single-cell suspensions were prepared immediately before analysis by flow cytometry as previously described^40^. In brief, postnatal and adult hearts were minced and digested in 2.5 mg/mL collagenase D (Roche), 0.25 mg/mL DNase I (Roche) and 0.05% Trypsin-EDTA solution in RPMI (Sigma) incubated for 45 minutes at 37°C accompanied with constant pipetting for mechanical separation. Digested samples were passed through a 70-μm Nylon cell strainer, washed and suspended in Hank’s Balanced Salt Solution (HBSS, Gibco) with 3% FBS and 0.03 mM EDTA (FACS buffer) for staining.

### Flow cytometry of cultured cells

Murine ES-derived CPC were analyzed for the presence of appropriate markers on designated days of mesodermal and cardiac differentiation with the use of an ARIA II Analyzer (BD Biosciences) and FACSDiva 7.0 software as previously described^39^. In brief, cultured cells were treated with 0.05% trypsin/EDTA (Gibco) for 5 min at 37°C under 5% CO_2_. Cells were labelled with the following antibodies: anti-human/mouse GFRA2 Polyclonal Goat IgG (R&D Systems), rat monoclonal anti-PDGFR beta antibody conjugated with PE (Abcam, APB5), donkey polyclonal anti-goat IgG Alexa 405 conjugated with UV (Abcam), rat monoclonal IgG2b anti-mouse KDR-Alexa647 conjugated with APC (BioLegend). 7-AAD (BioLegend, Cat no. 420404) was used as a viability marker. Sca-1 (Biolegend Cat no. 108127), c-Kit (Biolegend, Cat no. 105813), CD31 (Biolegend, Cat no. 102524). Flow Cytometry data analysis was performed using FlowJo^™^ V10.

### FACS-sorted cell culture conditions

Acquired sorted cells were harvested in 50% FBS in PBS and centrifuged for 20 min at 4°C. After centrifuging, cells were seeded onto 96-well laminin-coated plates and cultured in 20%FBS/DMEM (Gibco) with penicillin-streptomycin (Gibco). Medium was changed every 3 days and cells were fixed on designated days for further analysis.

### Cell culture immunofluorescence staining

Cultured cells were fixed with pre-warmed 4% paraformaldehyde (PFA) for 10 min at room temperature. Fixed cells were washed three times for 5 min in PBS, and then nonspecific antibody binding sites were blocked with blocking buffer 1% BSA/ 0.2% Triton X-100 in PBS for 30 min at RT. After that cells were incubated with primary antibodies in blocking buffer overnight at 4°C. Primary antibodies: Cardiac troponin T (cTnT, 1/100, mouse monoclonal, Abcam), alpha smooth muscle actin (aSMA) (1/100, rabbit polyclonal, Abcam), CD31 (1/25, Rabbit polyclonal, Abcam), anti-GFP FITC-conjugated (1/100, goat polyclonal, Abcam).

After rinsing 3 times for 5 min with PBS, cells were incubated with secondary antibodies for 1h at RT: Alexa Fluor 555 Goat anti-mouse IgG, Alexa Fluor 647 Goat anti-rabbit IgG. Again, cells were washed three times for 5 min in PBS and then mounted with DAPI mounting medium (Fluoroshield with DAPI, Sigma-Aldrich). Images were acquired with an inverted Leica DMIRE2 microscope and a Hamamatsu Camera ORCA Flash 4.0 LT.

### Heart tissue sectioning and staining

Adult murine hearts were isolated and fixed in 4% PFA in PBS and embedded in paraffin, sectioned transversely at 5-μm thickness and loaded onto slides. The sections underwent deparaffinization with xylene and a decreasing ethanol gradient and were routinely stained with Haematoxylin and Eosin (H&E). Masson’s Trichrome staining was used in paraffin sections to identify collagen fibers in 1-month damaged murine hearts. Upon deparaffinization, stainings were used as followed: Hematoxyline Harris 1 min, Red of Mallory 3min (Fuchsin acid, Sigma), Molybdophosphoric acid 1% 2min (dodeca-Molybdophosphoric acid, Vyzas), Methyl blue 1 min (Sigma). Between different stains slides were washed with dH2O. Lastly, slides were dehydrated with: 100% EtOH (1st) 1 min, 2nd EtOH 2min, acidified EtOH 2min, 1st Xylene 4min, 2nd Xylene 4min.

For immunohistochemistry, acquired adult murine hearts (and parts of) were perfused and fixed with 4% PFA in PBS, for 2h at 4°C. Then, they were transferred in 30% sucrose in PBS, at 4°C overnight. The next day they were embedded in OCT compound (VWR) and cryosections were prepared at 16 μm thickness. For immunohistochemistry, cryosections were post-fixed with pre-warmed 4% PFA in PBS for 15 min, and then rinsed in PBS to remove OCT. Non-specific antibody binding sites were blocked with 2% BSA (fraction V)/ 10% FBS/ 0.05% Tween 20 in PBS, at room temperature (RT) for 1.5h. Samples were incubated with primary antibodies in the blocking buffer at 4°C overnight. Primary antibodies used were: MF20 (mouse monoclonal, 1/100, Developmental Biology Hybridoma Bank), GFP (chicken polyclonal, 1/1000, Abcam), Tbx5 (rabbit polyclonal, 1/100, Sigma), GFRA2 (chicken polyclonal, 1/500, Antibodies-online, ABIN1450225), Connexin 43 (cat no C6219-.2ML, rabbit polyclonal, 1/2000, Sigma), α-actinin (A7811, clone EA-53, mouse monoclonal, 1/500, Sigma).

After washing 3 times with 0.5% Triton X-100 in PBS (PBST), the samples were incubated in the secondary antibody in the blocking buffer for 1 hour at RT. Secondary antibodies were used as followed: Alexa Fluor 555 Goat Anti-mouse IgG, Alexa Fluor 647 Goat Anti-rabbit IgG, Alexa Fluor 488-conjugated Goat Anti-chicken. After washing 3 times with PBST sections were mounted with DAPI. Fluorescent images were captured using an upright Leica DMRA2 fluorescence microscope and a Hamamatsu ORCA-Flash 4.0 V2.

### Reverse transcription and quantitative real-time PCR analysis

Hearts were collected on days 2, 4 and 7 after injury, and total RNA was extracted using TRIzol Reagent (Sigma-Aldrich, T9424) and chloroform (AppliChem). For each specimen 500 ng of total RNA was reversed transcribed into cDNA using PrimeScript RT reagent kit (TaKaRa RR037a) and oligo dT primers according to the manufacturer’s protocol. Prior to cDNA synthesis, samples were subjected to TURBO^™^ DNase (Invitrogen) treatment for 1h at 37°C and 10 min at 75°C. Quantitate PCR was conducted using KAPA SYBR FAST Master Mix (Sigma-Aldrich, KK4611) on a Roche Lightcycler 96 (Roche Life Science). Cycling conditions were as follows: 2 min at 50°C and 10 min at 95°C (Pre-incubation) followed by two-step PCR for 40 cycles of 15s at 95°C and 60s at 60°C. Expression levels were calculated using the comparative CT method and calculated 2^-ΔΔCt^ values are presented. Values for specific genes were normalized to *GAPDH* as a constitutively expressed internal control. Primers used are;

*eYFP:* F: 5′-ACGTAAACGGCCACAAGTTC-3′ R: 5′-AAGTCGTGCTGCTTCATGTG-3′
*Tbx5:* F: 5′-CTCCCAGCAAGTCTCCATCA-3′ R: 5′-GGCCAGTCACCTTCACTTTG-3′
*Gapdh:* F: 5′-AGGTCGGTGTGAACGGATTTG-3′ R: 5′-TGTAGACCATGTAGTTGAGGTCA-3′

### Statistics

Flow cytometric data are presented as mean ± SD. Statistical analysis on FACS data was performed using ANOVA T-test using Mann-Whitney or Bonferroni post-hoc test, where appropriate (p<0.05). Statistical analyses were calculated using GraphPad Prism 5. Single-cell RNA-seq data statistical analysis was initially performed using p-value (<0.05) and EdgeR after Benjamini and Hochberg^41^ for obtaining DE genes (DEGs). Downstream analysis involved False Discovery Analysis (FDR) based on Benjamini and Hochberg^41^. All statistical analyses were performed using GraphPad Prism, with the threshold for significance set at *P* < 0.05.

## RESULTS

### *In vitro* mESC-derived CPC follow a similar cardiac developmental program to *in vivo* early embryo CPCs

We set to establish an *in vitro* mesodermal/cardiac differentiation pipeline that would pinpoint an optimal developmental window where CPCs could be examined and collected for further studies (**Figure 1**). Based on a previously established CPCs differentiation system^39^, we were able to enrich for FHF and SHF CPCs populations based on their surface expression profile. Murine BAC *Tbx5^CreERT2^/Rosa26R^eYFP/eYFP^* ESCs were subjected to cardiac differentiation, while 4-hydroxytamoxifen (4OH-TAM) was added from day 4 onwards (**Figure 1A**). When CPCs were allowed to differentiate for up to 12 days *in vitro,* under non-FBS defined culture conditions, YFP (Tbx5-lineage traced) expression was observed in cTnT^+^ cells but never into endothelial nor smooth muscle cells (**Figure 2B**). In suboptimal differentiation conditions, there was an absence of YFP^+^ with a subsequent decrease in cTnT^+^ cells, in total (**Supplementary Figure 1B**).

**Figure 1.**
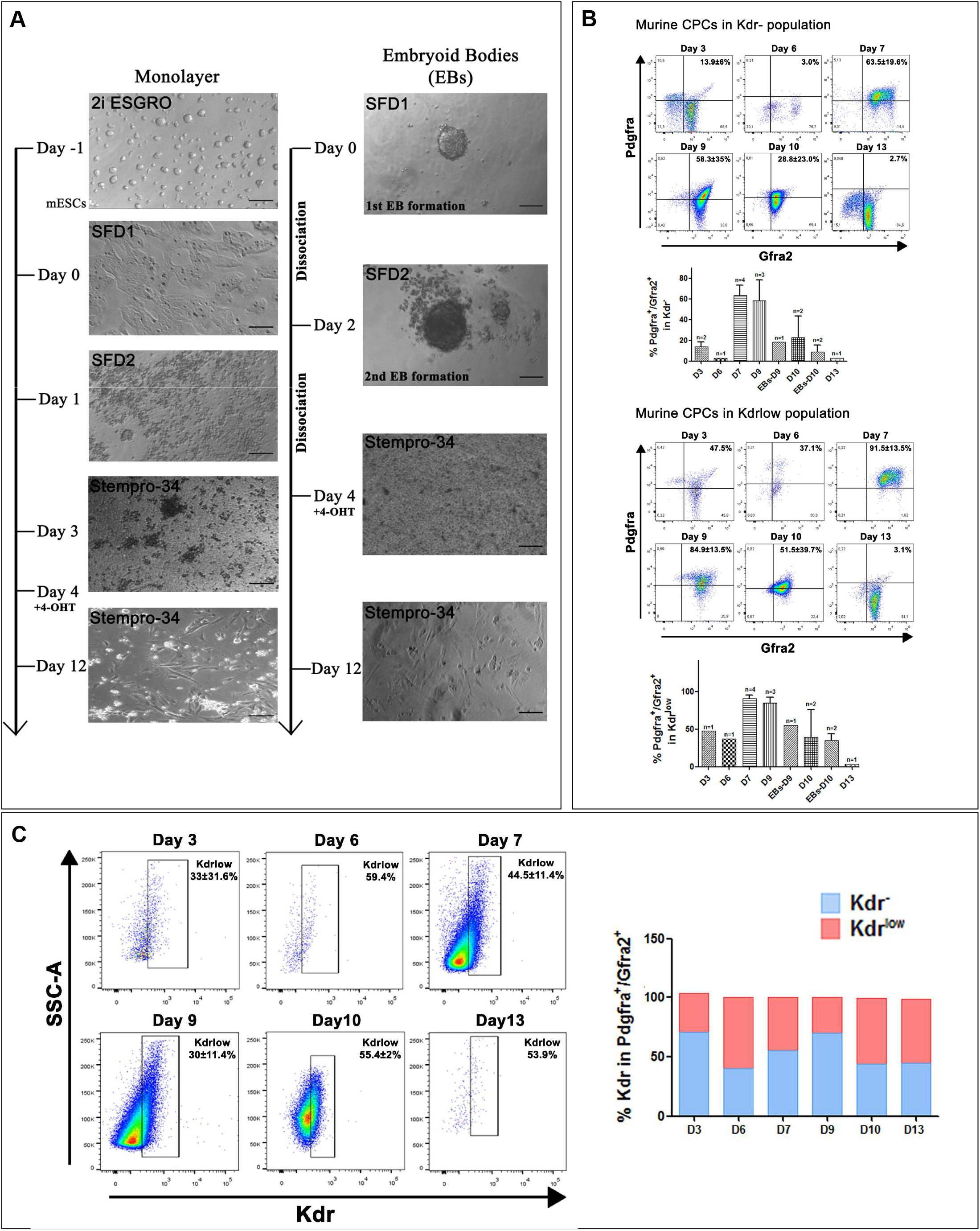
Enrichment and optimisation of in vitro mESC-derived CPC enrichment. **(A)** Representative microphotographs depicting stages of differentiation of murine ground state *Tbx5^Cre^*;*R26R^eYFP/eYFP^* mESC cultured under specified cardiomyocyte differentiation conditions, either as monolayers or as embryoid bodies. **(B)** Flow cytometric analysis using three surface markers Pdgfra, Kdr and Gfra2 on different days of cardiomyocyte differentiation indicates an CPC enrichment window between days 7 and 9. **(C)** The expression of Kdr is dynamic and defines two waves of CPC in the current differentiation protocol. N=1-4.

**Figure 2.**
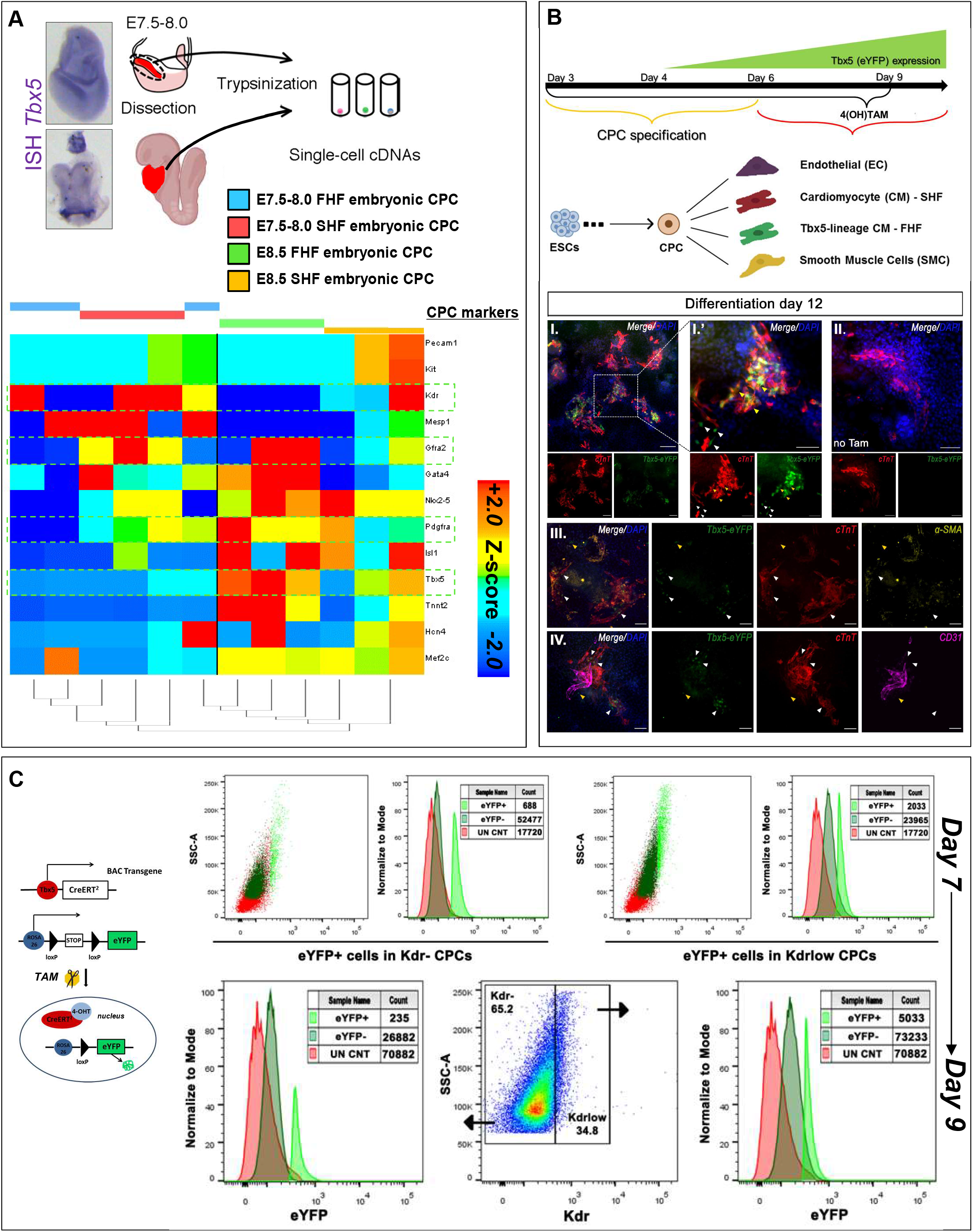
Tbx5 expression is enriched in CPC-derived cardiomyocytes. **(A)** Tbx5 expression in E7.5-E8.0 and E8.5 whole embryos according to mRNA *in situ* hybridisation. Single-cell RNA-seq analysis of cardiac cells from E7.5-E8.0 and E8.5 embryos for known cardiac progenitor cell genes. **(B)** Representative microphotographs of 4OH-TAM-induced YFP (Tbx5-lineage tracing) expression (green) on cardiomyocytes (cTnT, red), endothelial cells (CD31, purple) and smooth muscle cells (*α*-SMA, yellow) that have differentiated from mESC-derived CPC on day 12. N=5. **(C)** Flow cytometric analysis indicates that BAC *Tbx5^Cre^*;*R26R^eYFP/eYFP^* insert can lineage trace Tbx5-expressing cells within the CPC population. N=9. Scale bar = 10 μm.

Flow cytometric analysis indicated that Pdgfra^+^/Gfra2^+^/Kdr^low/+^ CPCs were enriched between days 7 and 9 (**Figure 1B, Supplementary Figure 1**). Interestingly, in our *in vitro* culture system, Kdr surface marker pinpointed to two potential “waves” of Pdgfra^+^/Gfra2^+^ CPCs (**Figure 1C**). To address this finding, we revisited previously published data from our lab concerning single-cell RNA-seq obtained from *in vivo* cardiac CPCs on different early embryonic stages^23^ (**Figure 2**). Gene expression metaanalysis indicated that embryonic day 8.5 (E8.5) CPC showed a decreased *Kdr* expression, when compared to E7.5 CPCs, in an inversely proportional manner to *Tbx5* expression, *in vivo* (**Figure 2A**). The expression of YFP after 4OH-TAM administration, indicated that both Kdr^low/+^ and Kdr^-^CPC (Pdgfra^+^/Gfra2^+^) subpopulations possessed YFP^+^ CPCs, yet only the Kdr^low/+^/Gfra2^+^/Pdgfra^+^/YFP^+^CPC population increased from days 7 to 9 (**Figure 2C**).

These data indicate that the *in vitro* mESC-derived CPCs’ surface marker expression profile follows a gene expression profile similar to that of *in vivo* CPCs involved in early cardiac embryonic development, with Tbx5^+^ CPCs to promote a unipotent CM-like fate both *in vivo* and *in vitro*.

### Reactivation of the TF Tbx5 in the adult injured mammalian heart

Using a BAC *Tbx5^CreERT2/CreERT2^/Rosa26R^eYFP/eYFP^* transgene^23^, we employed two adult heart injury murine models, a myocardial infraction (MI) of ischaemia/reperfusion (I/R), and a chemical injury model using ISO intraperitoneal (i.p.) injections to promote cardiomyocyte lesions in the cardiac muscle^42^ (**Figure 3A**). Cardiac injury was confirmed using Haematoxylin and Eosin as well as Masson’s trichrome staining (**Supplementary Figure 2**). Immunohistochemical analysis indicated that Tbx5^+^/YFP^+^ cells were present in the injured ventricles four days after injury, while Tbx5^-^YFP^+^ cells were present in injured ventricles on days 7 and 30 post-injury; no Tbx5^+^/YFP^+^ were detected in uninjured ventricles, as expected^43^ (**Figure 3B and Supplementary Figures 3 and 4**). No eYFP^+^cells were readily observed in uninjured adult LV (**Figure 3C**). To characterise these YFP^+^ further, adult injured heart immunohistochemistry indicated their co-expression with cardiogenic precursor markers such as Tbx5, Nkx2-5, but not Isl1. In addition, YFP^+^ did not co-localise with classical smooth muscle cell (*α*-SMA) nor immune cell markers (CD45), while CD31 and C-kit protein expression was observed in both YFP^+^ and YFP^-^ cells (**Figure 4A**). By investigating the whole heart after injury, the presence of YFP^+^ CM was primarily observed around lesion sites (**Figure 4B**). Confocal imaging indicated a disorganisation of those YFP^+^ cells’ sarcomere structure and gap junctions following injury, peaking around days 4-7, while on some YFP^+^ CM, gap junctions were partially restored by day 30 (**Figure 4C**).

**Figure 3.**
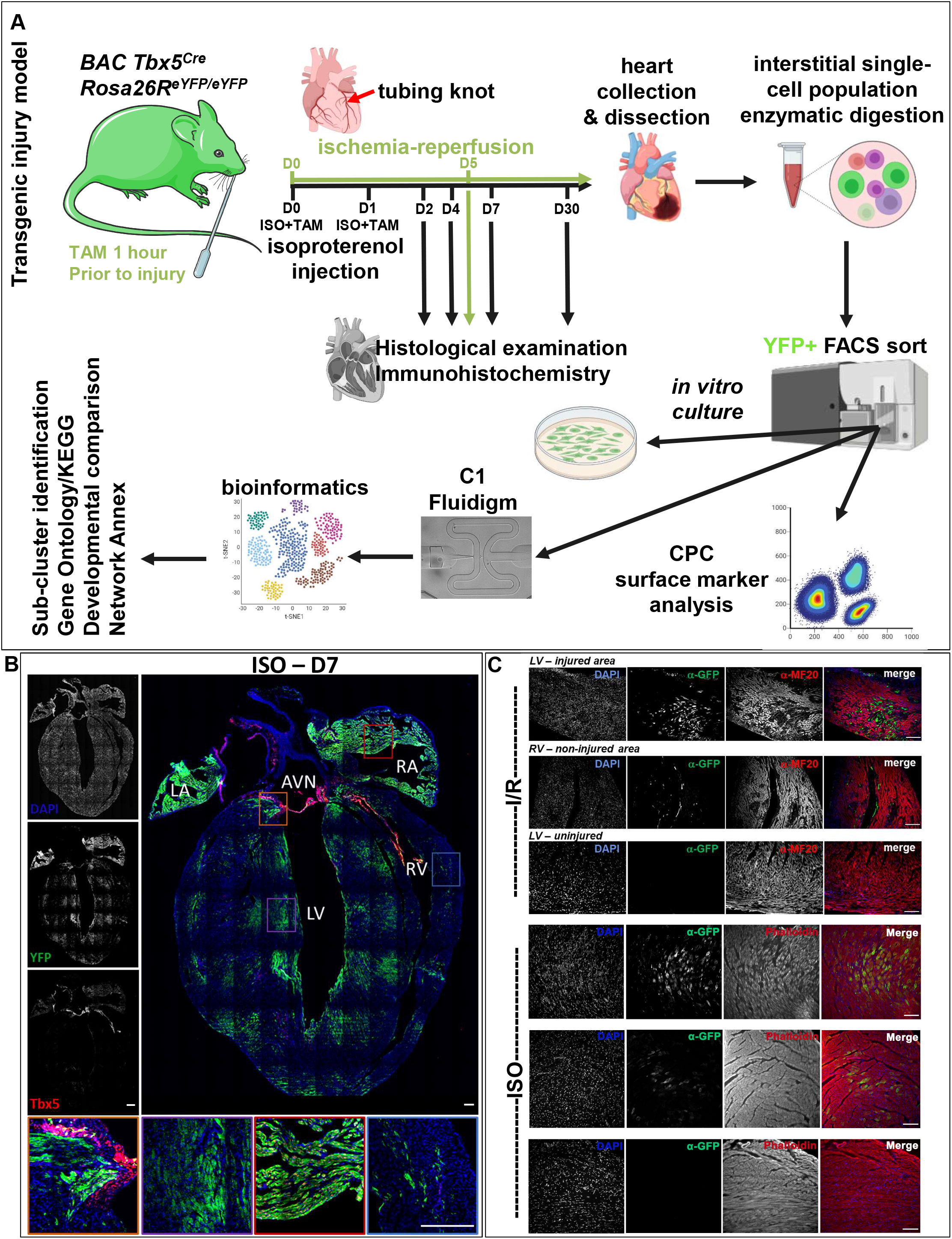
Tbx5 re-activation in the adult murine heart. **(A)** Schematic representation of the experimental pipeline for Tbx5-lineage (YFP^+^) tracing cell analysis in the adult injured murine heart. **(B)** A collage of an ISO-injured adult heart on D7 after injury. Key-LV=Left Ventricle, RV=Right Ventricle, RA=Right Atrium, LA=Left Atrium, AVN=Atrioventricular Node, SAN=Sinoatrial Node. Higher magnification inserts indicating the localisation of YFP^+^ cells in sites of ventricular injury, while YFP^+^Tbx5^+^ and Tbx5^+^YFP^-^ where only located in the atria and the nodes. **(C)** Representative microphotographs of adult hearts 5 days after I/R and 7 days after ISO injury, indicating YFP^+^ cells (green) and cardiomyocytes [MF20 or phalloidin/F-actin, red]. N=2-3 hearts per condition and n>6 sections per heart examined. Scale bar = 10 μm.

**Figure 4.**
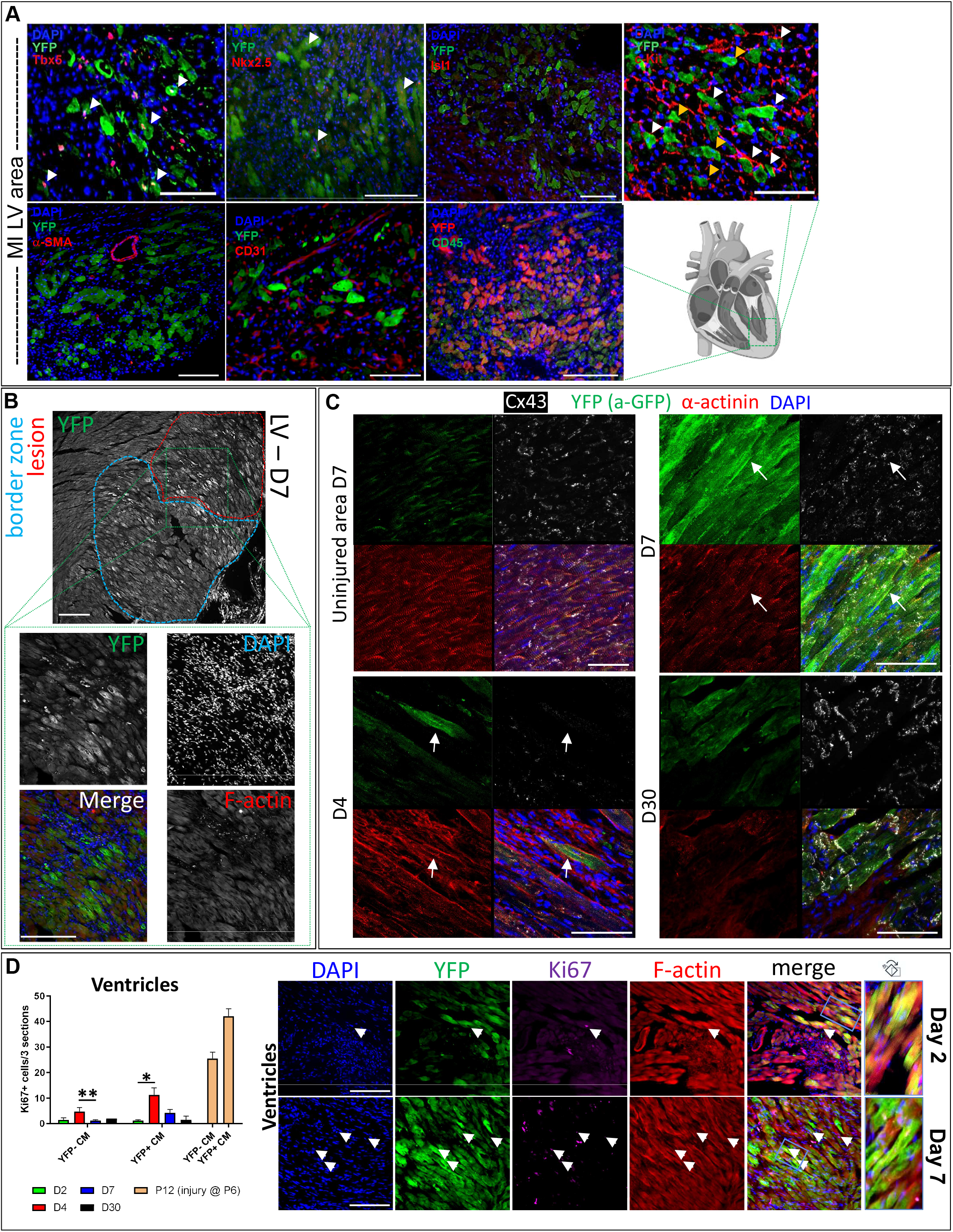
In situ characterization of YFP^+^ cells upon heart injury. **(A)** Representative microphotographs of immunohistochemistry performed in adult injured heart cryosections on day 5 after I/R. **(B)** Representative microphotographs of a lesion site where increased YFP^+^ CMs are present in the lesion and border zones. Higher magnification inserts indicate YFP^+^F-actin^+^ cardiac cells with a CM morphology located in and around the lesion site. **(C)** Representative microphotographs of lesions in adult injured hearts 2, 4, 7 and 30 days post injury; YFP^+^-expressing cells (green), cardiomyocyte sarcomeres (α-actinin, red), gap-junction protein Connexin-43 (Cx43, white) and nuclear dye (DAPI, blue). Scale bars = as indicated. **(D)** XY graph depicting Ki67^+^ cells per section in YFP^+^ and YFP^-^ CM in adult injured hearts, 2, 4 and 7 days post-injury, as well as 7 days post-injury in P6 pups. YFP^+^-expressing cells (green), cardiomyocytes (F-actin/Phalloidin, red), cycling cells (Ki67, magenta) and nuclear dye (DAPI, blue). Arrowheads indicative of co-localisation of YFP^+^-expressing cells (green), cardiomyocytes (F-actin/Phalloidin, red), cycling cells (Ki67, magenta) and nuclear dye (DAPI, blue). Blue insert magnifies on a representative YFP^+^ cell that has altered sarcomere striation designated by F-actin, when compared to an adjacent YFP^+^F-actin^+^ CM. *D2 vs D4 p=0.0019, **D4 vs D7 p=0.009583. N=1-3 hearts per time-point and n>6 sections per heart examined. Scale bar = 10 μm.

Immunohistochemical analysis at different time points indicated that some YFP^+^ CMs showed a low, yet persistent expression of the cell cycle protein Ki67, when compared to YFP^-^ CMs (**Figure 4D**). These results are in line with recent findings of Ki67 expression in adult CM populations^44^.

Flow cytometric analysis was performed in order to collect eYFP^+^ single cells from 2, 4, 5 and 7 days post injury. The expression of YFP and Tbx5 transcripts in the atria and ventricles of the adult injured heart were also confirmed by qPCR (**Figures 5A, B** and **Supplementary Figure 5A**). The eYFP protein expression was validated in cardiac ventricular cell populations in the injured hearts, with a peak YFP expression 7 days after injury (**Figure 5B**).

**Figure 5.**
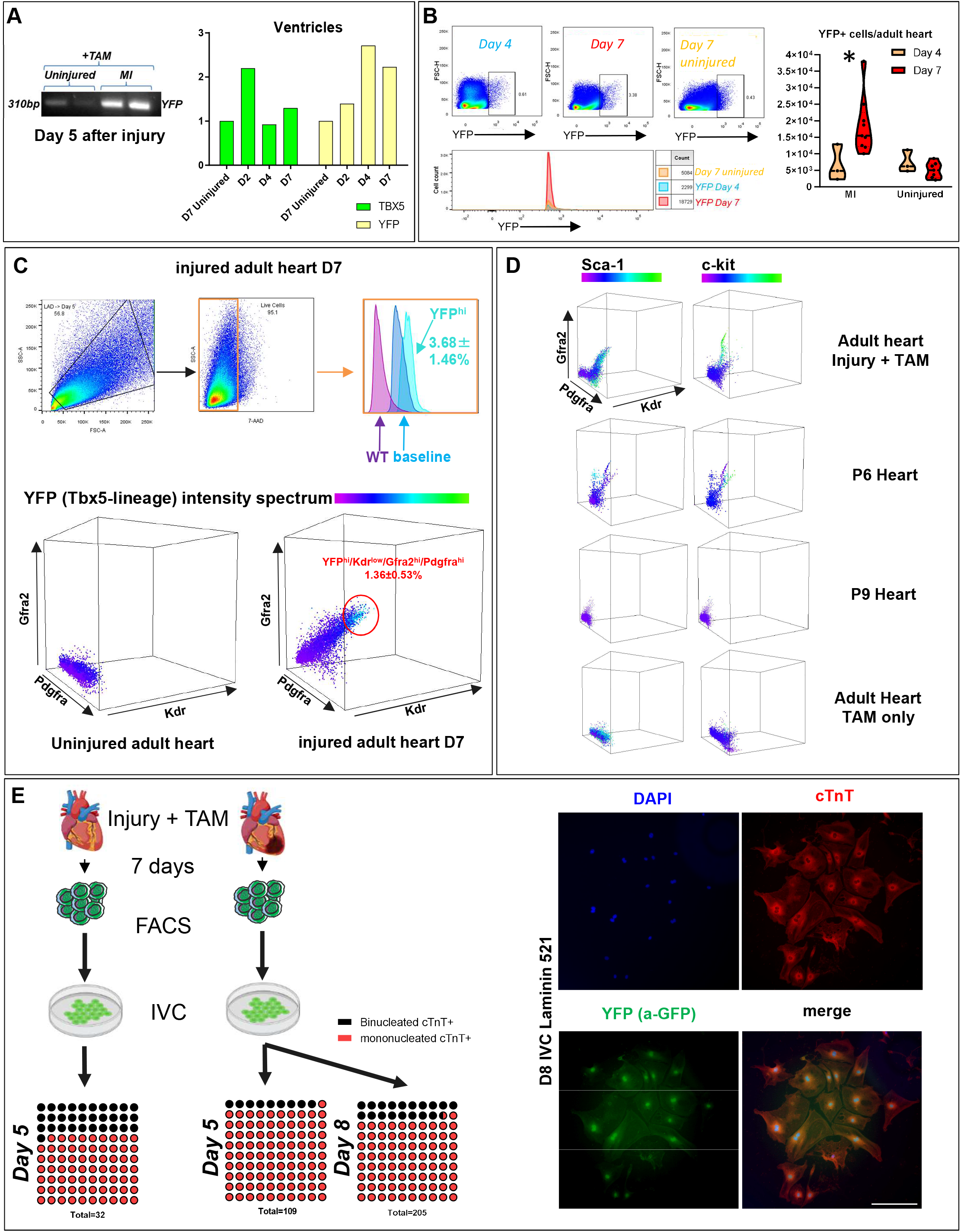
CPC surface analysis of YFP^+^ cells. **(A)** Real-time PCR analysis of *Yfp* and *Tbx5* transcripts in the ventricles of the adult heart in different time-points. **(B)** Flow cytometry acquisition of YFP^+^ cells from different time-points following cardiac injury. N=3-6 hearts. **(C, D)** Representative two- and threedimensional graphs from flow cytometric analysis of Pdgfra^+^Kdr^low/+^Gfra2^+^ adult, postnatal days 6 and 9 heart and their co-expression with YFP (Tbx5-tracing), Sca-1 and c-Kit. N=9. Key-CP; cardiomyocyte precursor, CM; cardiomyocyte, EC; endothelial cell. **(E)** YFP^+^ cells were collected 7 days after chemical injury and cultured in CM differentiating conditions for 5 and 8 days. Only YFP^+^ from the injured heart were able to differentiate into CM-like cells. Measurement of YFP^+^/cTnT^+^ mononucleated and binucleated cells 5 and 8 days *in vitro* culture. N=3 independent cultures.

The finding that an embryonic TF such as Tbx5 is being reactivated in the adult injured heart, led us to investigate whether the well-documented cell-surface embryonic CPC markers^45,46^, may be able to tag an adult cardiac cell subpopulation. FACS analysis was performed on cells collected from adult lungs and hearts (treated with Tamoxifen only) or injured adult hearts, as well as hearts derived from postnatal (P) days 6 and 9. Results showed that the YFP^+^ cells are a part of a wider Kdr^low/+^/Gfra2^+^/Pdgfra^+^ adult cardiac cell population (**Figure 5C and Supplementary Figure 5B**). Our data also showed that a Kdr^low/+^/Gfra2^+^/Pdgfra^+^ CPC subpopulation was detected only in cells derived from P6 whole heart tissue, but were near-absent in P9 hearts, neither in uninjured adult murine hearts nor lungs (**Figure 5D and Supplementary Figure 5C**). Of note, Sca-1 was detected, but could not be designated in Kdr^low/+^/Gfra2^+^/Pdgfra^+^/YFP^+/-^ cells only, while c-kit was not detected in our adult CPCs, yet present in P6 CPC (Kdr^low/+^/Gfra2^+^/Pdgfra^+^), as shown previously^47^.

Recent studies have demonstrated that CM polyploidy is relevant to their regenerative capacity, with polynucleation acting as a barrier against regeneration^48^. Collected YFP^+^ adult cells were cultured *in vitro* 7 days after injury, for 5 and 8 days (**Figure 5E**). It was observed an increased frequency of mononucleated YFP^+^cTnT^+^ cell population in cells collected from the whole injured heart, in comparison to the whole uninjured heart (31.7±5.6% uninjured vs 90.8±11.1% injured on Day 5).

Taken together, these data indicate that a Tbx5-expressing potentially precursor CM population is activated upon myocardial injury, in the adult mammalian heart.

### Injury-induced YFP^+^ CM precursors may support a reparative potential of the adult CM ventricular tissue

In order to assess the pathophysiology of Tbx5-expressing heart cells, we employed an mESC Tbx5-KO primary cell line, from which we attempted to obtain CPCs using a defined differentiation medium ^23,39^. We observed a delay in the early Kdr^low/+^/Gfra2^+^/Pdgfra^+^ CPC formation (Day 7). This postponement was compensated later on (Day 10) (**Supplementary Figure 6A**). To assess the potential pathophysiology of Tbx5-expressing CM *in vivo,* upon injury we created a tamoxifen-induced *Tbx5^CreERT2/+^/Rosa26R^eYFP/+^/Rosa26R^iDTR/+^* transgene, where upon TAM administration, Tbx5-expressing cells will undergo apoptosis. It was observed that adult mice that received one dose of ISO and TAM had a 50% increased lethality, when compared to mice that only received TAM (**Supplementary Figure 6B**). Histological examination 4 days-post injury showed severe LV and atrial injury in *Tbx5^CreERT2/+^/Rosa26R^eYFP/+^/Rosa26R^iDTR/+^* mice that received ISO+TAM, while administration of TAM showed atrial injury only along with reduced ventricular damage, when compared to ISO+TAM (**Supplementary Figure 6C**). The cause of death was possibly due to reduced lung branching after the massive loss of alveolar Tbx5-expressing cells causing bronchiectasis (data not shown). These findings place Tbx5 at a pivotal point for cardiac ventricular repair.

### The transcriptome of adult mammalian injury-induced YFP^+^ cells resembles that of developmentally earlier cardiac cells

By employing single-cell RNA-seq (scRNA-seq) deep sequencing analysis on 116 YFP^+^-sorted cells and comparing them to Pdgfra^+^ uninjured interstitial cells, it was possible to confirm their distinct expression profile and reveal least two major YFP^+^ cell sub-clusters (**Figure 6A**). Heatmap analysis on the FACS markers employed in this study and reference cardiac fibroblast genes^49^ confirmed their enrichment for *Tbx5*, *Gfra2*, *Kdr and Pdgfra*, and their underrepresentation, respectively in YFP^+^ cells, in relation to Pdgfra^+^ interstitial cardiac cells (**Figure 6B**). In order to biologically identify and separate the two most prominent YFP^+^ cell sub-clusters we further statistically analysed their DEGs (**Figure 6C**). Gene Ontology (GO) and KEGG enrichment analysis revealed differences in oxidative phosphorylation, cardiac muscle development and morphogenesis, thermogenesis and hormonal responses (diabetic cardiomyopathy)^50^ as well as signalling related to central nervous system input (**Figure 7A**).

**Figure 6.**
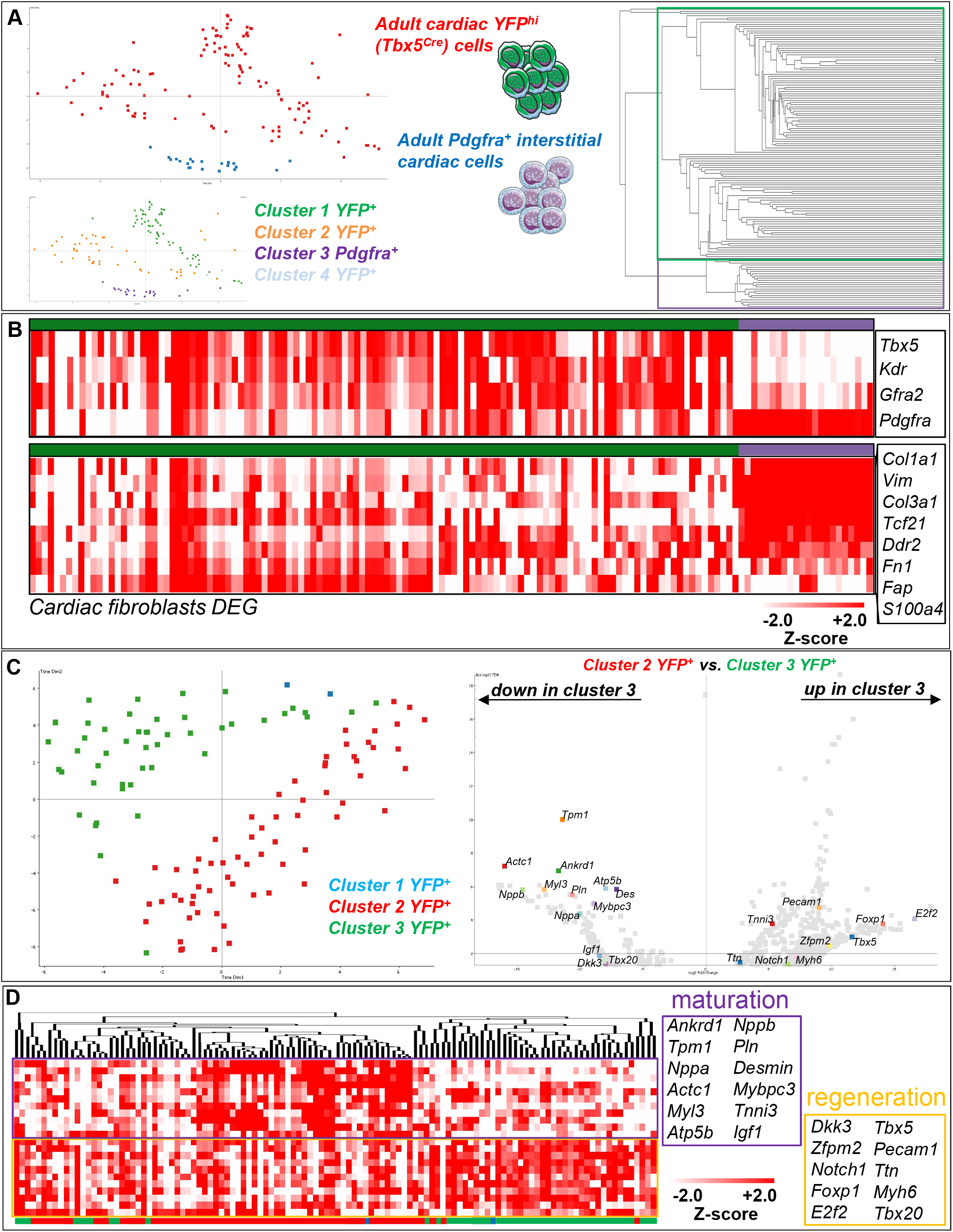
The transcriptome of YFP^+^ cell sub-clusters resembles that of CM precursors. **(A)** t-SNE dimensionality analysis and data store tree of EdgeR p<0.05 after Benjamimi and Hochberg correction (8,749 DEGs) of YFP^+^ (green) and Pdgfra^+^ (blue) interstitial adult heart cells. YFP^+^ cells could be divided into at least three sub-clusters. Single cells examined were obtained from 3 adult injured D7 hearts. **(B)** Heatmaps depicting expression of *Tbx5*, *Kdr*, *Gfra2 and Pdgfra* and cardiac fibroblast DEGs in YFP^+^ (green) and uninjured Pdgfra^+^ (purple) adult heart cells. **(C)** t-SNE dimensionality, and volcano plot of DEGs between the two major YFP^+^ sub-clusters (1,091 DEGs). **(D)** Heatmap clustering analysis of CM-relevant genes within the DEG list that showed prominent expression differences between the two YFP^+^ sub-clusters.

**Figure 7.**
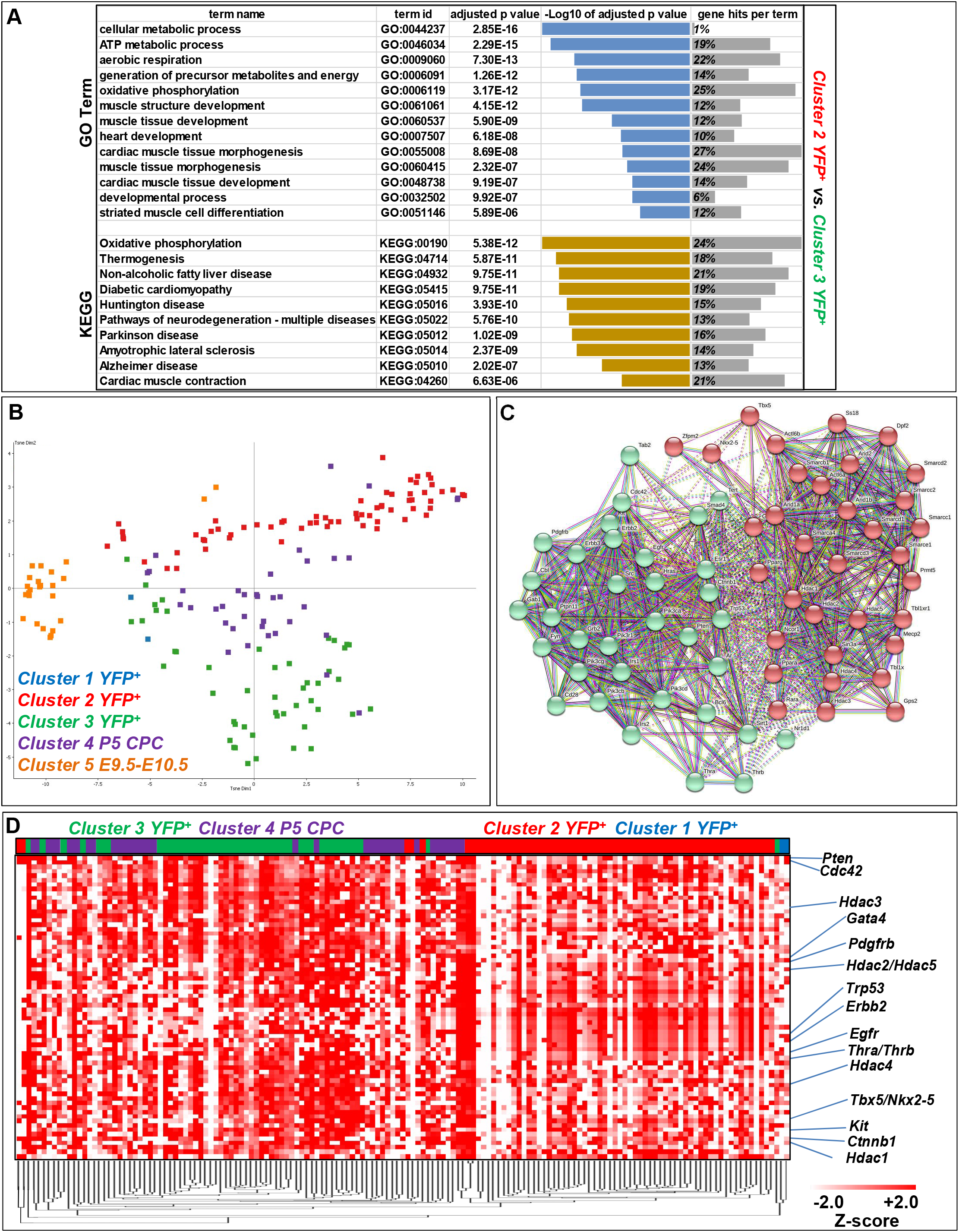
Developmental comparison of YFP^+^ CM. **(A)** Highlighted GO Biological Process and KEGG enrichment terms are shown. p<0.025, FDR<0.05 (Benjamimi and Hochberg correction). **(B)** t-SNE dimensionality analysis of all probes between YFP^+^ cell sub-clusters 1-3 (blue, red, green), P5 CPC (purple) and embryonic heart cells collected from E9.5 and E10.5 (orange). **(C)** *Kmeans* STRING transcriptional network diagram of *mus musculus* Tbx5-related and Thrα/β proteins interactions. **(D)** Heatmap clustering analysis of STRING gene expression in YFP^+^ sub-clusters 1-3 (blue, red, green) and P5 CPC (purple) cells.

The transcription factor Tbx5 has been shown to be expressed in embryonic cardiac cells that possess CM precursor properties, which can be faithfully recapitulated using our *Tbx5^CreERT2/CreERT2^*;*Rosa26R^eYFP/eYFP^* transgene, in *ex vivo* settings (**Supplementary Figure 7A**)^51^. To investigate the adult YFP^+^ cardiac cell population in-depth, a roadmap of single-cell RNA-seq transcriptomes was created from published single-cell RNA-seq *in vivo* E9.5-E10.5 cardiac progenitors^33^, as well as from CPC deriving from postnatal day 5, where cardiac regeneration is still achievable in mice (**Figure 7B**). Clustering analysis underscored that adult YFP^+^ cells from the injured hearts were transcriptomically relevant to postnatal cardiac CPC populations, while more distant from early embryonic cardiac progenitors. *Kmeans* STRING clustering analysis indicates that the Tbx5 transcriptional network is directly linked to thyroid hormonal responses (**Figure 7C**). Based on recent studies that involve thermogenesis and thyroid hormonal regulation in CM regeneration^50,52,53^, as well as our *in silico* meta-analysis data for thyroid hormone receptors binding to Tbx5 (data not shown), we interrogated the transcriptional profile of P5 CPC and YFP^+^ CMs in relation to the two major protein clusters, Tbx5 and Thyroid Hormone Receptors alpha and beta (Thrα/Thrβ) (**Figure 7D**). Heatmap clustering analysis indicated a similar expression pattern of Tbx5-related and Thrα/β-related genes in YFP^+^ CM cluster 3 and P5 CPC cells, when compared to YFP^+^ CM cluster 2.

## DISCUSSION

The developing mammalian murine heart, initially, shares common progenitors with mesodermal progenitors of the cranial and paraxial mesoderm^54^, which have shown to give rise to muscle with regenerative capabilities^55,56^; the adult heart mammalian muscle lacks this property. Even if the injured heart has any substantial regenerative capacity, this is lost after the first week of age, in mice^57,58^. In the recent years, several studies have identified resident cells, which have been implicated in cardiomyocyte regeneration, yet these evidence have been heavily debated. Eventually, it has been concluded that *de novo* cardiac stem cells are not present in the adult mammalian heart^59–62^.

The accumulated knowledge from embryonic cardiac development and CPCs has not been readily utilised in the adult regenerative field, with a few exceptions (Nkx2-5, Isl1) [For a review, please see ^63^]. In the current study, we employed a well-characterised transgenic mouse model capable of spatiotemporally lineage-tracing Tbx5^+^ cells during cardiac embryonic development^23^ and now, in adult mammalian hearts. The fact that this is a BAC insert, allows for investigating the spatiotemporal expression of Tbx5 in the adult heart, without inducing a cardiac phenotype^23^. Importantly, our transgene is capable of lineage-tracing mostly ventricular Tbx5-expressing cardiac cells, which may be important for ventricle-specific repair/regeneration, in-line with recent published basic research and preclinical data^64–66^. In our study, Tbx5 re-activated CMs were located close to the lesion areas, but a few were also observed in non-injured areas of the adult heart. Recently, a major single-cell RNA-seq study using frozen human ischemic heart biopsies, designated *TBX5* overexpression in CM subpopulations as an important protective mechanism (pre-print server https://doi.org/10.1101/2021.06.23.449672).

In humans, TBX5 is expressed in both the embryonic and adult four-chambered heart, in yet unidentified cardiac cell populations^67^. In the adult mouse, Tbx5 has been shown to be primarily expressed in the atria but not in the ventricles, in the absence of injury, which we also observed^43^. We show that during chemical and I/R heart injuries, a CM-like ventricular subpopulation transiently re-activates Tbx5. Albeit present a month after injury, our quantification analysis did not show the formation of new *in vivo* YFP^+^-derived CM (data not shown). In our experimental setting, we observed YFP^+^ CM with a disrupted sarcomere and gap junctions structures peaking between 4 and 7 days after injury, with gap junctions, being partially restored a month after injury in some of those YFP^+^ CM. *In vitro,* YFP^+^ cells were capable of expressing CM markers but not smooth muscle or endothelial cell-like cells. Thus, it cannot be excluded that a low-level regeneration capability exists in the adult mammalian heart, albeit unable to compensate for the massive CM loss that occurs during a major heart incidence.

One question that arises is whether a specific pre-designated CM subpopulation is primed for initiating CM replenishment, and/or repair, during injury, or what observed is actually a stochastic spatiotemporal event? We answer this question by underscoring the significance of Tbx5 in the adult heart; by omitting Tbx5-expressing cells *via* induced cell death, myocardial injury is aggravated. It is currently debatable whether an elusive regenerative mechanism in mammals is promoted by the presence of an endogenous pre-existing CM-like precursor population or a CM that directly re-enters the cell cycle^62^. Yet, these two possibilities are not mutually exclusive; they could imply that during myocardial insult, a subpopulation of CMs undergoes dedifferentiation (i.e. transiently becoming CM precursors) through a process called epimorphosis, in order to proliferate and provide with new CMs. Indeed, a dedifferentiated regressive CM population that has regained a primitive phenotype, has been recently reported in adult murine and human injured hearts^68,69^. It has been documented that non-cardiomyocyte cell populations are actively proliferating in homeostatic postnatal^70^ and injured adult murine hearts^71^, showing no signs of apoptosis^72^. This has been also confirmed in our immunohistochemistry data, showing that proliferative interstitial cardiac cell populations were also present. Our results are indicative of an adult Tbx5-overexpressing ventricular cardiac cell that fits the profile of a potentially dedifferentiated CM-like precursor that fails to proliferate/re-differentiate, even if some cell cycle activators are expressed; they may form a septation-like zone around the injury site acting as guidepost cells^73^. In concert with our findings, a *Tbx5* mRNA expression burst has been documented in border-zone viable CMs^68^. Interestingly, the authors of the same study identified that the cardiac-specific *natriuretic peptide precursor type A* (*Nppa*) gene is highly expressed in those border-zone CMs; its expression is activated by the binding of Tbx5 and Nkx2-5 TFs in the *Nppa* promoter region^74^.

We reasoned that the Tbx5-lineage-tagged YFP^+^ adult cardiac population could be a potential cardiac precursor candidate, and for this we employed knowledge gained from embryonic cardiac development, which can define and characterise that candidate^46^. The aforementioned CPC status of Gfra2^+^Pdgfra^+^Kdr^low/+^YFP^+^ cells was validated *in vitro*, in *Tbx5^CreERT2/CreERT2^*;*Rosa26R^eYFP/eYFP^* mESC-derived CPCs prior to proceeding with our *in situ* investigation. We show here that an adult cardiac cell population exists in the injured mammalian heart that resembles the more well-defined embryonic cardiac precursors and postnatal precursors, in the surface marker expression levels and mRNA transcriptome, respectively. The presence of this triple surface-marker signature population observed only in the injured adult murine heart was also confirmed upon evaluation of metadata from previous scRNA-seq cardiac studies (analysis not shown)^75–77^. Of interest, not all of the adult Gfra2^+^Pdgfra^+^Kdr^low/+^ cardiac cells, expressed YFP. Yet, over 70% of the YFP^+^ cells did express Gfra2 and Pdgfra, while their Kdr expression profile was dynamic, similar to our mESC-derived CPC *in vitro* and embryonic data.

Interestingly, between P1 and P7, which is the only reported postnatal regenerative window of the murine heart, Gfra2^+^Pdgfra^+^Kdr^low/+^ cells were present in the absence of an insult. It remains to be seen if these cells are responsible for the aforementioned remuscularisation window, which has been recently shown to be affected by metabolic cues^78^. We therefore, set to explore through single-cell transcriptomic analysis how the adult YFP^+^ ventricular cells resemble early postnatal (P5) Gfra2^+^Pdgfra^+^Kdr^low/+^ and embryonic cardiac cells. Our findings reveal an adult cell population that transcriptomically resembles that of postnatal CM and less that of embryonic cardiac cells, a finding that is in line with adult limb regeneration studies conducted in amphibians^79^.

The apparent lack of an efficient *in vivo* regeneration upon mammalian adult cardiac injury may be attributed into two main distinct conditions; (i) an idle population capable of producing new CM, and (ii) a microenvironment that deters CM regeneration favouring resilience, thus increasing the chances of survival of the organism as a whole in the short-term (i.e. inducing sustainable fibrosis to avoid cardiac rupture)^80^. We report here that the first condition could be met, while the second one has been shown to be indeed a major hurdle for heart regeneration^50^. As such, endogenous CM-like precursors are present, yet unable to reach their true/full intrinsic regenerative potential.

Our studies raise the question of whether there is a distinct developmental origin of those adult Tbx5-expressing CM precursors of the LV, similar to what has been observed in the primitive streak^81^. Recently, Zhang *et al.* through an extensive *Mesp1*^+^-lineage tracing and scRNA-seq analyses, revealed that the FHF possess at least two early distinct cardiomyogenic progenitors, from which, a subset of the LV CMs, derived from Tbx5^+^ CPCs^82^. Here, we report that the YFP^+^ CM-like population is potentially responsive to CNS-derived signals, hinting towards a cardiac neural crest origin, as shown recently in the zebrafish^83^.

Bae *et al*. reported that malonate injections during MI, were capable of enhancing heart regeneration in adult mice^78^. The target CM subpopulation that drove the regeneration was not examined. It would be intriguing to assess whether the Tbx5^+^ lineage-traced population identified in our studies, is one of the plastic CM subpopulations affected by malonate and/or thyroxine, as also reported recently^50^.

One limitation of this study was the inability to collect YFP^+^ immediately after cardiac injury; this was due to the intracellular expression delay of the YFP protein (48 hours after Tbx5 expression), which has been noted in our model and others’ (^23^ and references within). Therefore, although it is possible to assume that a similar cardiac regeneration window exists in the adult (as in neonates), where Tbx5 is transiently expressed, we were unable to collect a reliable number of valid YFP^+^ cells for further lineage-tracing analysis.

Another limitation is the absence of a reference point where adult mammalian CM regeneration is apparent. This hinders our ability to report that the Tbx5-expressing CM precursor cell population identified in the adult injured mammalian ventricles will eventually replenish the lost CMs. To overcome this, future studies will be also focusing on neonatal murine cardiac injuries where a natural CM regeneration exists. Although this has been elegantly shown in adult non-mammalian organisms^21^, definitive experiments in mammals will allow us to confirm the potential of Tbx5-expressing CM precursors and definitively show that the latter are part of an idle mammalian cardiac regenerative program that is standing by.

In conclusion, this study reveals and characterises a Tbx5-expressing ventricular CM-like precursor compartment identified following cardiac injury. As such, trending regenerative approaches can be tailored to target and trace the aforementioned cardiac cell population, which when found within a positive microenvironment, may induce adult CM regeneration.

## ACKNOWLEDGEMENTS

We thank Dr. Anastasia Apostolidou for handling the FACS facility at the BRFAA. Many thanks to Prof Kenta Yashiro, for being an excellent mentor and for the revision of this manuscript prior to its submission. We also thank Mr Dimitrios Valakos for his help with *in silico* meta-analysis.

## REFERENCES

1 Virani, S. S. et al. Heart Disease and Stroke Statistics-2020 Update: A Report From the American Heart Association. Circulation 141, e139–e596, doi:10.1161/CIR.0000000000000757 (2020).

2 Evans, S. M., Yelon, D., Conlon, F. L. & Kirby, M. L. Myocardial lineage development. Circ Res 107, 1428–1444, doi:10.1161/CIRCRESAHA.110.227405 (2010).

3 Rana, M. S., Christoffels, V. M. & Moorman, A. F. A molecular and genetic outline of cardiac morphogenesis. Acta Physiol (Oxf) 207, 588–615, doi:10.1111/apha.12061 (2013).

4 Downs, K. M. & Davies, T. Staging of gastrulating mouse embryos by morphological landmarks in the dissecting microscope. Development 118, 1255–1266 (1993).

5 Tam, P. P. & Behringer, R. R. Mouse gastrulation: the formation of a mammalian body plan. Mech Dev 68, 3–25 (1997).

6 Buckingham, M., Meilhac, S. & Zaffran, S. Building the mammalian heart from two sources of myocardial cells. Nat Rev Genet 6, 826–835, doi:10.1038/nrg1710 (2005).

7 Ma, Q., Zhou, B. & Pu, W. T. Reassessment of Isl1 and Nkx2-5 cardiac fate maps using a Gata4-based reporter of Cre activity. Dev Biol 323, 98–104, doi:10.1016/j.ydbio.2008.08.013 (2008).

8 Hsieh, P. C. et al. Evidence from a genetic fate-mapping study that stem cells refresh adult mammalian cardiomyocytes after injury. Nat Med 13, 970–974, doi:10.1038/nm1618 (2007).

9 Bergmann, O. et al. Evidence for cardiomyocyte renewal in humans. Science 324, 98–102, doi:10.1126/science.1164680 (2009).

10 Mollova, M. et al. Cardiomyocyte proliferation contributes to heart growth in young humans. Proc Natl Acad Sci U S A 110, 1446–1451, doi:10.1073/pnas.1214608110 (2013).

11 Ali, S. R. et al. Existing cardiomyocytes generate cardiomyocytes at a low rate after birth in mice. Proc Natl Acad Sci U S A 111, 8850–8855, doi:10.1073/pnas.1408233111 (2014).

12 Malliaras, K. et al. Cardiomyocyte proliferation and progenitor cell recruitment underlie therapeutic regeneration after myocardial infarction in the adult mouse heart. EMBO Mol Med 5, 191–209, doi:10.1002/emmm.201201737 (2013).

13 Senyo, S. E. et al. Mammalian heart renewal by pre-existing cardiomyocytes. Nature 493, 433–436, doi:10.1038/nature11682 (2013).

14 Steimle, J. D. & Moskowitz, I. P. TBX5: A Key Regulator of Heart Development. Curr Top Dev Biol 122, 195–221, doi:10.1016/bs.ctdb.2016.08.008 (2017).

15 Bruneau, B. G. et al. Chamber-specific cardiac expression of Tbx5 and heart defects in Holt-Oram syndrome. Dev Biol 211, 100–108, doi:10.1006/dbio.1999.9298 (1999).

16 Jia, Y., Chang, Y., Guo, Z. & Li, H. Transcription factor Tbx5 promotes cardiomyogenic differentiation of cardiac fibroblasts treated with 5-azacytidine. J Cell Biochem, doi:10.1002/jcb.28885 (2019).

17 Kathiriya, I. S. et al. Modeling Human TBX5 Haploinsufficiency Predicts Regulatory Networks for Congenital Heart Disease. Dev Cell, doi:10.1016/j.devcel.2020.11.020 (2020).

18 Inagawa, K. et al. Induction of Cardiomyocyte-like Cells in Infarct Hearts by Gene Transfer of Gata4, Mef2c, and Tbx5. Circ Res, doi:CIRCRESAHA.112.271148 [pii] 10.1161/CIRCRESAHA.112.271148 (2012).

19 McDonnell, T. J. & Oberpriller, J. O. The response of the atrium to direct mechanical wounding in the adult heart of the newt, Notophthalmus viridescens. An electron-microscopic and autoradiographic study. Cell Tissue Res 235, 583–592, doi:10.1007/BF00226956 (1984).

20 Grajevskaja, V., Camerota, D., Bellipanni, G., Balciuniene, J. & Balciunas, D. Analysis of a conditional gene trap reveals that tbx5a is required for heart regeneration in zebrafish. PLoS One 13, e0197293, doi:10.1371/journal.pone.0197293 (2018).

21 Sanchez-Iranzo, H. et al. Tbx5a lineage tracing shows cardiomyocyte plasticity during zebrafish heart regeneration. Nat Commun 9, 428, doi:10.1038/s41467-017-02650-6 (2018).

22 Wu, C. C. et al. Spatially Resolved Genome-wide Transcriptional Profiling Identifies BMP Signaling as Essential Regulator of Zebrafish Cardiomyocyte Regeneration. Dev Cell 36, 36–49, doi:10.1016/j.devcel.2015.12.010 (2016).

23 Kokkinopoulos, I. et al. Single-Cell Expression Profiling Reveals a Dynamic State of Cardiac Precursor Cells in the Early Mouse Embryo. PLoS One 10, e0140831, doi:10.1371/journal.pone.0140831 (2015).

24 Smemo, S. et al. Regulatory variation in a TBX5 enhancer leads to isolated congenital heart disease. Hum Mol Genet 21, 3255–3263, doi:10.1093/hmg/dds165 (2012).

25 Uehara, M., Yashiro, K., Takaoka, K., Yamamoto, M. & Hamada, H. Removal of maternal retinoic acid by embryonic CYP26 is required for correct Nodal expression during early embryonic patterning. Genes Dev 23, 1689–1698, doi:10.1101/gad.1776209 (2009).

26 Buch, T. et al. A Cre-inducible diphtheria toxin receptor mediates cell lineage ablation after toxin administration. Nat Methods 2, 419–426, doi:10.1038/nmeth762 (2005).

27 Lindsey, M. L. et al. Guidelines for in vivo mouse models of myocardial infarction. Am J Physiol Heart Circ Physiol 321, H1056–H1073, doi:10.1152/ajpheart.00459.2021 (2021).

28 Chatzianastasiou, A. et al. Cardioprotection by H2S Donors: Nitric Oxide-Dependent and Independent Mechanisms. J Pharmacol Exp Ther 358, 431–440, doi:10.1124/jpet.116.235119 (2016).

29 Brooks, W. W. & Conrad, C. H. Isoproterenol-induced myocardial injury and diastolic dysfunction in mice: structural and functional correlates. Comp Med 59, 339–343 (2009).

30 Song, Y. et al. Cardiac ankyrin repeat protein attenuates cardiac hypertrophy by inhibition of ERK1/2 and TGF-beta signaling pathways. PLoS One 7, e50436, doi:10.1371/journal.pone.0050436 (2012).

31 Oudit, G. Y. et al. Phosphoinositide 3-kinase gamma-deficient mice are protected from isoproterenol-induced heart failure. Circulation 108, 2147–2152, doi:10.1161/01.CIR.0000091403.62293.2B (2003).

32 Miyao, N. et al. TBX5 R264K acts as a modifier to develop dilated cardiomyopathy in mice independently of T-box pathway. PLoS One 15, e0227393, doi:10.1371/journal.pone.0227393 (2020).

33 Li, G. et al. Transcriptomic Profiling Maps Anatomically Patterned Subpopulations among Single Embryonic Cardiac Cells. Dev Cell 39, 491–507, doi:10.1016/j.devcel.2016.10.014 (2016).

34 Dobin, A. et al. STAR: ultrafast universal RNA-seq aligner. Bioinformatics 29, 15–21, doi:10.1093/bioinformatics/bts635 (2013).

35 Li, H. et al. The Sequence Alignment/Map format and SAMtools. Bioinformatics 25, 2078–2079, doi:10.1093/bioinformatics/btp352 (2009).

36 Kokkinopoulos, I. et al. Patrolling human SLE haematopoietic progenitors demonstrate enhanced extramedullary colonisation; implications for peripheral tissue injury. Sci Rep 11, 15759, doi:10.1038/s41598-021-95224-y (2021).

37 Chen, J., Bardes, E. E., Aronow, B. J. & Jegga, A. G. ToppGene Suite for gene list enrichment analysis and candidate gene prioritization. Nucleic Acids Res 37, W305–311, doi:10.1093/nar/gkp427 (2009).

38 Bindea, G. et al. ClueGO: a Cytoscape plug-in to decipher functionally grouped gene ontology and pathway annotation networks. Bioinformatics 25, 1091–1093, doi:10.1093/bioinformatics/btp101 (2009).

39 Kokkinopoulos, I. et al. Cardiomyocyte Differentiation From Mouse Embryonic Stem Cells Using a Simple and Defined Protocol. Dev. Dyn. 245, 157–165 (2016).

40 Pinto, A. R. et al. Revisiting Cardiac Cellular Composition. Circ Res 118, 400–409, doi:10.1161/CIRCRESAHA.115.307778 (2016).

41 Benjamini, Y. H., Y. Controlling the False Discovery Rate: A Practical and Powerful Approach to Multiple Testing. Journal of the Royal Statistical Society. Series B (Methodological) 57, 289–300 (1995).

42 Cowled, P. & Fitridge, R. in Mechanisms of Vascular Disease: A Reference Book for Vascular Specialists (eds R. Fitridge & M. Thompson) (2011).

43 Arnolds, D. E. et al. TBX5 drives Scn5a expression to regulate cardiac conduction system function. J Clin Invest 122, 2509–2518, doi:10.1172/JCI62617 (2012).

44 Huang, W. et al. Loss of microRNA-128 promotes cardiomyocyte proliferation and heart regeneration. Nat Commun 9, 700, doi:10.1038/s41467-018-03019-z (2018).

45 Kattman, S. J., Huber, T. L. & Keller, G. M. Multipotent flk-1+ cardiovascular progenitor cells give rise to the cardiomyocyte, endothelial, and vascular smooth muscle lineages. Dev Cell 11, 723–732 (2006).

46 Ishida, H. et al. GFRA2 Identifies Cardiac Progenitors and Mediates Cardiomyocyte Differentiation in a RET-Independent Signaling Pathway. Cell Reports 16, 1026–1038 (2016).

47 Jesty, S. A. et al. c-kit+ precursors support postinfarction myogenesis in the neonatal, but not adult, heart. Proc Natl Acad Sci U S A 109, 13380–13385, doi:10.1073/pnas.1208114109 (2012).

48 Patterson, M. et al. Frequency of mononuclear diploid cardiomyocytes underlies natural variation in heart regeneration. Nat Genet 49, 1346–1353, doi:10.1038/ng.3929 (2017).

49 Ivey, M. J. & Tallquist, M. D. Defining the Cardiac Fibroblast. Circ. J. 80, 2269–2276 (2016).

50 Hirose, K. et al. Evidence for hormonal control of heart regenerative capacity during endothermy acquisition. Science 364, 184–188, doi:10.1126/science.aar2038 (2019).

51 Bruneau, B. G. Signaling and transcriptional networks in heart development and regeneration. Cold Spring Harb Perspect Biol 5, a008292, doi:10.1101/cshperspect.a008292 (2013).

52 Pantos, C. & Mourouzis, I. Thyroid hormone receptor alpha1 as a novel therapeutic target for tissue repair. Ann Transl Med 6, 254, doi:10.21037/atm.2018.06.12 (2018).

53 Pantos, C. & Mourouzis, I. Translating thyroid hormone effects into clinical practice: the relevance of thyroid hormone receptor alpha1 in cardiac repair. Heart Fail Rev 20, 273–282, doi:10.1007/s10741-014-9465-4 (2015).

54 Tyser, R. C. V. et al. Characterization of a common progenitor pool of the epicardium and myocardium. Science, doi:10.1126/science.abb2986 (2021).

55 De Micheli, A. J. et al. Single-Cell Analysis of the Muscle Stem Cell Hierarchy Identifies Heterotypic Communication Signals Involved in Skeletal Muscle Regeneration. Cell Rep 30, 3583–3595 e3585, doi:10.1016/j.celrep.2020.02.067 (2020).

56 Dell’Orso, S. et al. Single cell analysis of adult mouse skeletal muscle stem cells in homeostatic and regenerative conditions. Development 146, doi:10.1242/dev.174177 (2019).

57 Mahmoud, A. I. et al. Meis1 regulates postnatal cardiomyocyte cell cycle arrest. Nature 497, 249–253, doi:10.1038/nature12054 (2013).

58 Porrello, E. R. et al. Transient regenerative potential of the neonatal mouse heart. Science 331, 1078–1080, doi:10.1126/science.1200708 (2011).

59 Vicinanza, C. et al. Kit(cre) knock-in mice fail to fate-map cardiac stem cells. Nature 555, E1–E5, doi:10.1038/nature25771 (2018).

60 van Berlo, J. H. et al. c-kit+ cells minimally contribute cardiomyocytes to the heart. Nature 509, 337–341, doi:10.1038/nature13309 (2014).

61 Ellison, G. M. et al. Adult c-kit(pos) cardiac stem cells are necessary and sufficient for functional cardiac regeneration and repair. Cell 154, 827–842, doi:10.1016/j.cell.2013.07.039 (2013).

62 Li, Y. et al. Genetic Lineage Tracing of Nonmyocyte Population by Dual Recombinases. Circulation 138, 793–805, doi:10.1161/CIRCULATIONAHA.118.034250 (2018).

63 Le, T. & Chong, J. Cardiac progenitor cells for heart repair. Cell death discovery 2, 16052, doi:10.1038/cddiscovery.2016.52 (2016).

64 Jiang, L. et al. CRISPR activation of endogenous genes reprogramsfibroblasts into cardiovascular progenitorcells for myocardial infarction therapy. Mol Ther, doi:10.1016/j.ymthe.2021.10.015 (2021).

65 Isomi, M. et al. Overexpression of Gata4, Mef2c, and Tbx5 Generates Induced Cardiomyocytes Via Direct Reprogramming and Rare Fusion in the Heart. Circulation 143, 2123–2125, doi:10.1161/CIRCULATIONAHA.120.052799 (2021).

66 Rathjens, F. S. et al. Preclinical evidence for the therapeutic value of TBX5 normalization in arrhythmia control. Cardiovasc Res 117, 1908–1922, doi:10.1093/cvr/cvaa239 (2021).

67 Hatcher, C. J., Goldstein, M. M., Mah, C. S., Delia, C. S. & Basson, C. T. Identification and localization of TBX5 transcription factor during human cardiac morphogenesis. Dev Dyn 219, 90–95, doi:10.1002/1097-0177(200009)219:1<90::AID-DVDY1033>3.0.CO;2-L (2000).

68 van Duijvenboden, K. et al. Conserved NPPB+ Border Zone Switches From MEF2- to AP-1-Driven Gene Program. Circulation 140, 864–879, doi:10.1161/CIRCULATIONAHA.118.038944 (2019).

69 Zhang, Y. et al. Single-cell imaging and transcriptomic analyses of endogenous cardiomyocyte dedifferentiation and cycling. Cell discovery 5, 30, doi:10.1038/s41421-019-0095-9 (2019).

70 O’Meara, C. C. et al. Transcriptional reversion of cardiac myocyte fate during mammalian cardiac regeneration. Circ Res 116, 804–815, doi:10.1161/CIRCRESAHA.116.304269 (2015).

71 Kretzschmar, K. et al. Profiling proliferative cells and their progeny in damaged murine hearts. Proc Natl Acad Sci U S A, doi:10.1073/pnas.1805829115 (2018).

72 Dispersyn, G. D. et al. Dissociation of cardiomyocyte apoptosis and dedifferentiation in infarct border zones. Eur Heart J 23, 849–857, doi:10.1053/euhj.2001.2963 (2002).

73 Waldron, L. et al. The Cardiac TBX5 Interactome Reveals a Chromatin Remodeling Network Essential for Cardiac Septation. Dev Cell 36, 262–275, doi:10.1016/j.devcel.2016.01.009 (2016).

74 Hiroi, Y. et al. Tbx5 associates with Nkx2-5 and synergistically promotes cardiomyocyte differentiation. Nat Genet 28, 276–280, doi:10.1038/90123 (2001).

75 McLellan, M. A. et al. High-Resolution Transcriptomic Profiling of the Heart During Chronic Stress Reveals Cellular Drivers of Cardiac Fibrosis and Hypertrophy. Circulation 142, 1448–1463, doi:10.1161/CIRCULATIONAHA.119.045115 (2020).

76 Farbehi, N. et al. Single-cell expression profiling reveals dynamic flux of cardiac stromal, vascular and immune cells in health and injury. eLife 8, doi:10.7554/eLife.43882 (2019).

77 Skelly, D. A. et al. Single-Cell Transcriptional Profiling Reveals Cellular Diversity and Intercommunication in the Mouse Heart. Cell Rep 22, 600–610, doi:10.1016/j.celrep.2017.12.072 (2018).

78 Bae, J. et al. Malonate Promotes Adult Cardiomyocyte Proliferation and Heart Regeneration. Circulation 143, 1973–1986, doi:10.1161/CIRCULATIONAHA.120.049952 (2021).

79 Lin, T. Y. et al. Fibroblast dedifferentiation as a determinant of successful regeneration. Dev Cell 56, 1541–1551 e1546, doi:10.1016/j.devcel.2021.04.016 (2021).

80 Notari, M. et al. The local microenvironment limits the regenerative potential of the mouse neonatal heart. Sci Adv 4, eaao5553, doi:10.1126/sciadv.aao5553 (2018).

81 Ivanovitch, K. et al. Ventricular, atrial, and outflow tract heart progenitors arise from spatially and molecularly distinct regions of the primitive streak. PLoS Biol 19, e3001200, doi:10.1371/journal.pbio.3001200 (2021).

82 Zhang, Q. et al. Unveiling Complexity and Multipotentiality of Early Heart Fields. Circ Res 129, 474–487, doi:10.1161/CIRCRESAHA.121.318943 (2021).

83 Tang, W., Martik, M. L., Li, Y. & Bronner, M. E. Cardiac neural crest contributes to cardiomyocytes in amniotes and heart regeneration in zebrafish. eLife 8, doi:10.7554/eLife.47929 (2019).

