## Supplementary Figures for "Return of the Tbx5; lineage-tracing reveals ventricular cardiomyocyte-like precursors in the injured adult mammalian heart"

**Supplementary Figure 1** **Siatra *et al.***

**A**

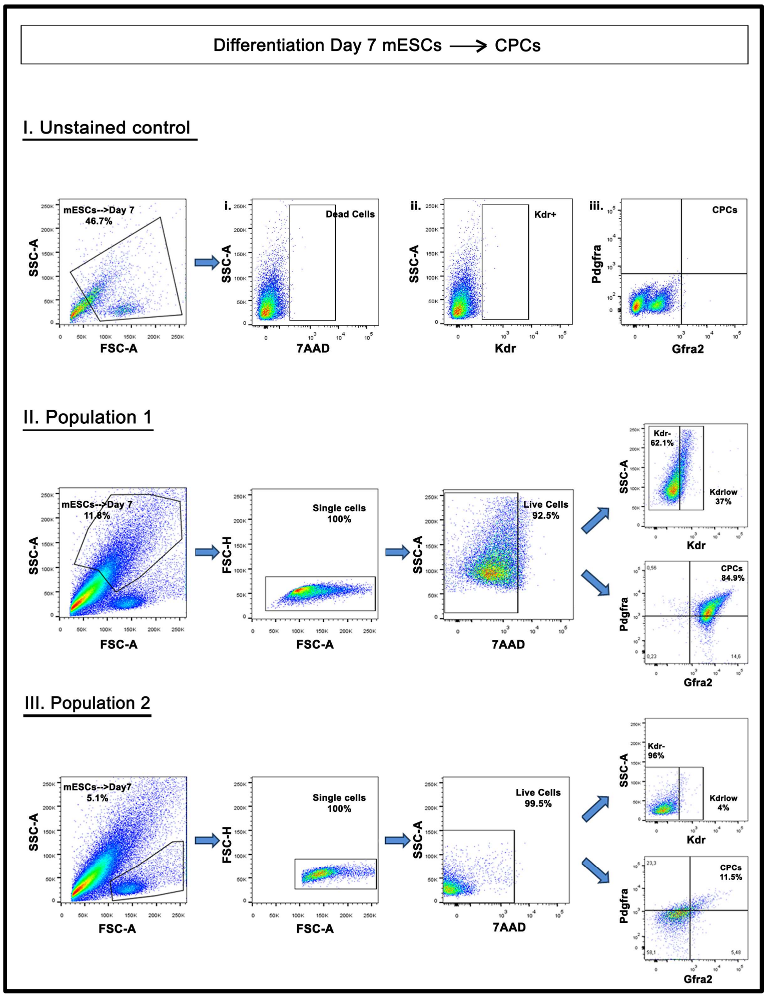


**B**

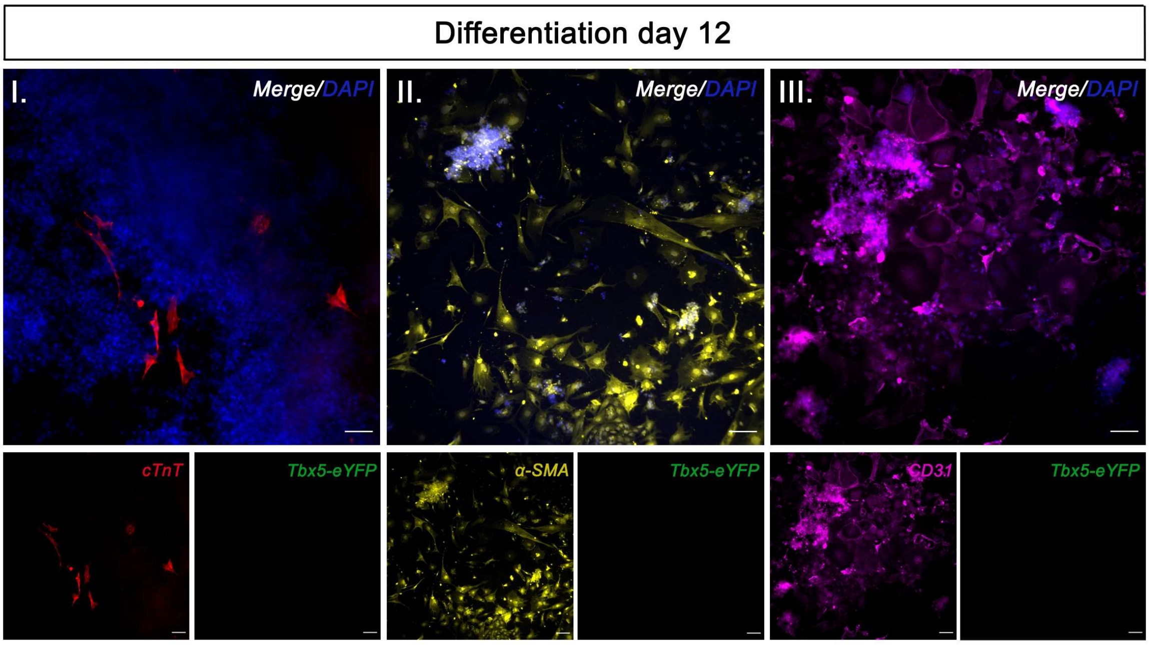


Supplementary Figure 1. (**A**) Gating strategy for acquiring triple positive cells from *in vitro* mESC in CM differentiation conditions, 7 days after 3i medium removal. (**B**) Immunocytochemical analysis of $Tbx5^{Cre}$;$R26R^{eYFP/eYFP}$ mESC-derived differentiated cells in suboptimal CM differentiation conditions, after 12 days *in vitro*. Scale bar = 100μm

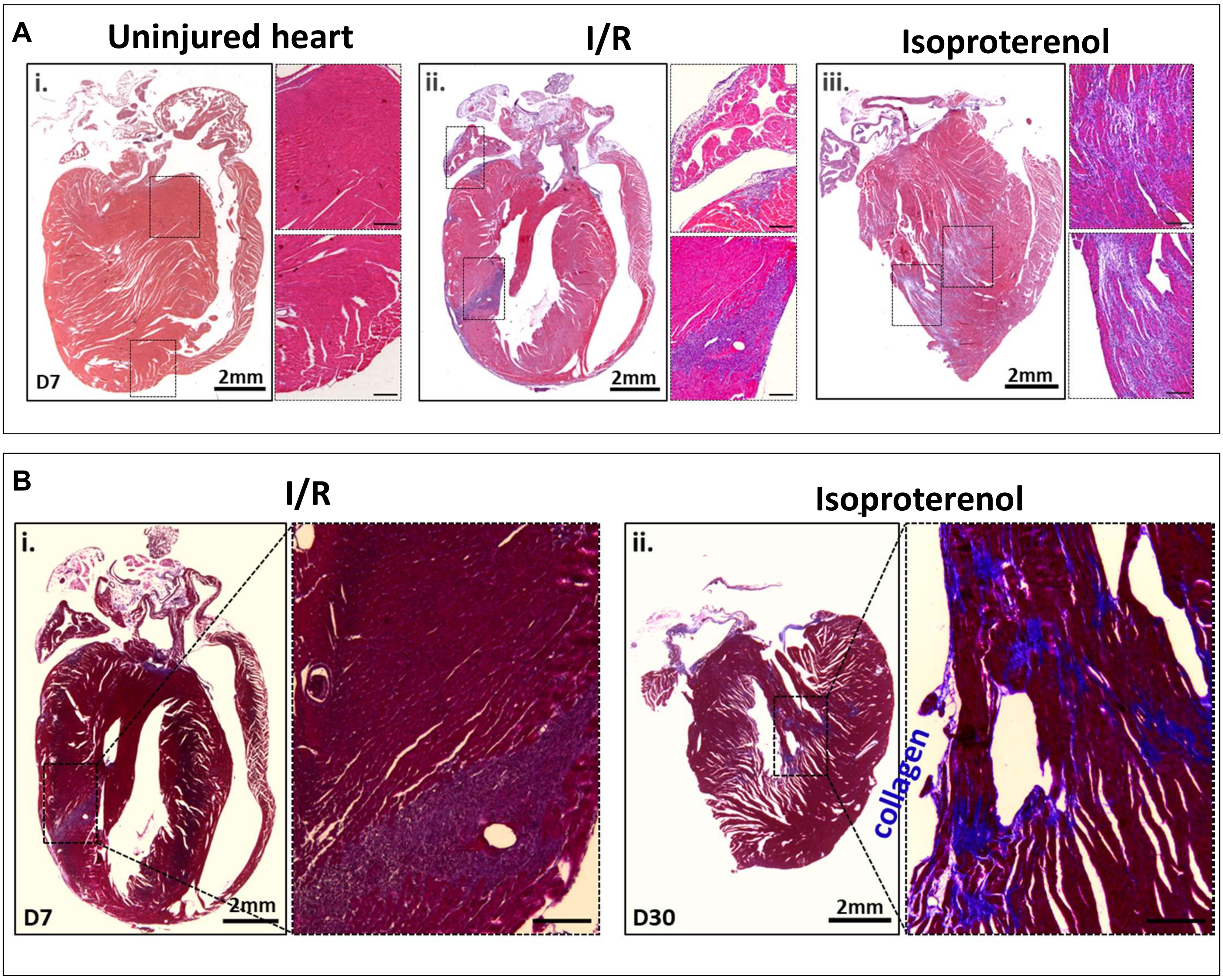


Supplementary Figure 2. (**A**) Haematoxylin and Eosin staining as well as whole heart photographs confirmed non-fatal, yet substantial, heart injury in the left ventricle, when compared to non-MI adult hearts. (**A**) Masson's staining, of collagen deposition at D7 and D30 after injury. N=1-3 per condition. Scale bar 100μm.

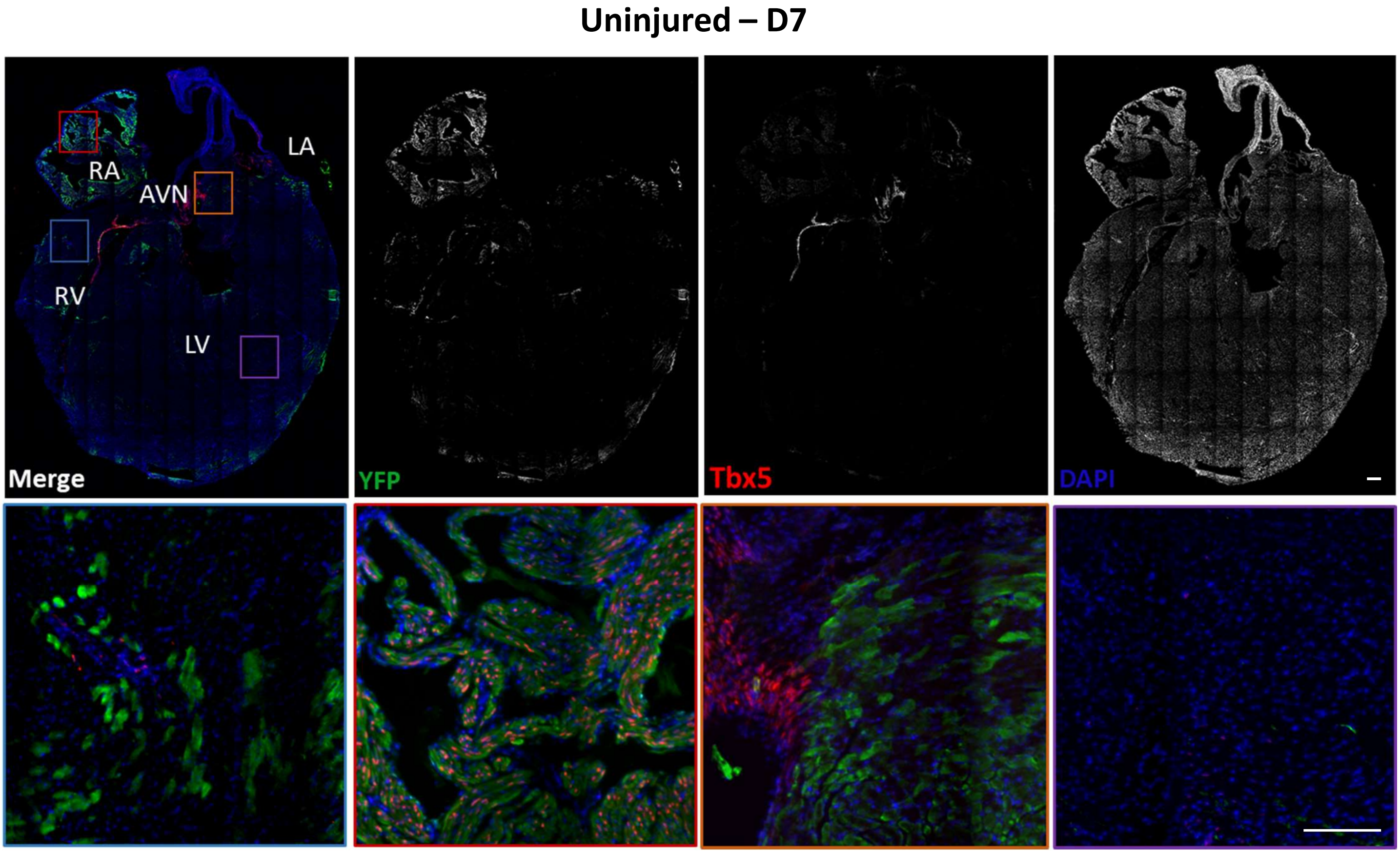


Supplementary Figure 3. (A) A collage of an uninjured adult heart 7 days after receiving TAM. Inserts indicating YFP⁺ and Tbx5⁺ are located in the Atria and AVN only. Key- LV=Left Ventricle, RV=Right Ventricle, RA=Right Atrium, LA=Left Atrium, AVN=Atrioventricular Node, N=2-3 hearts. Scale bar=100μm

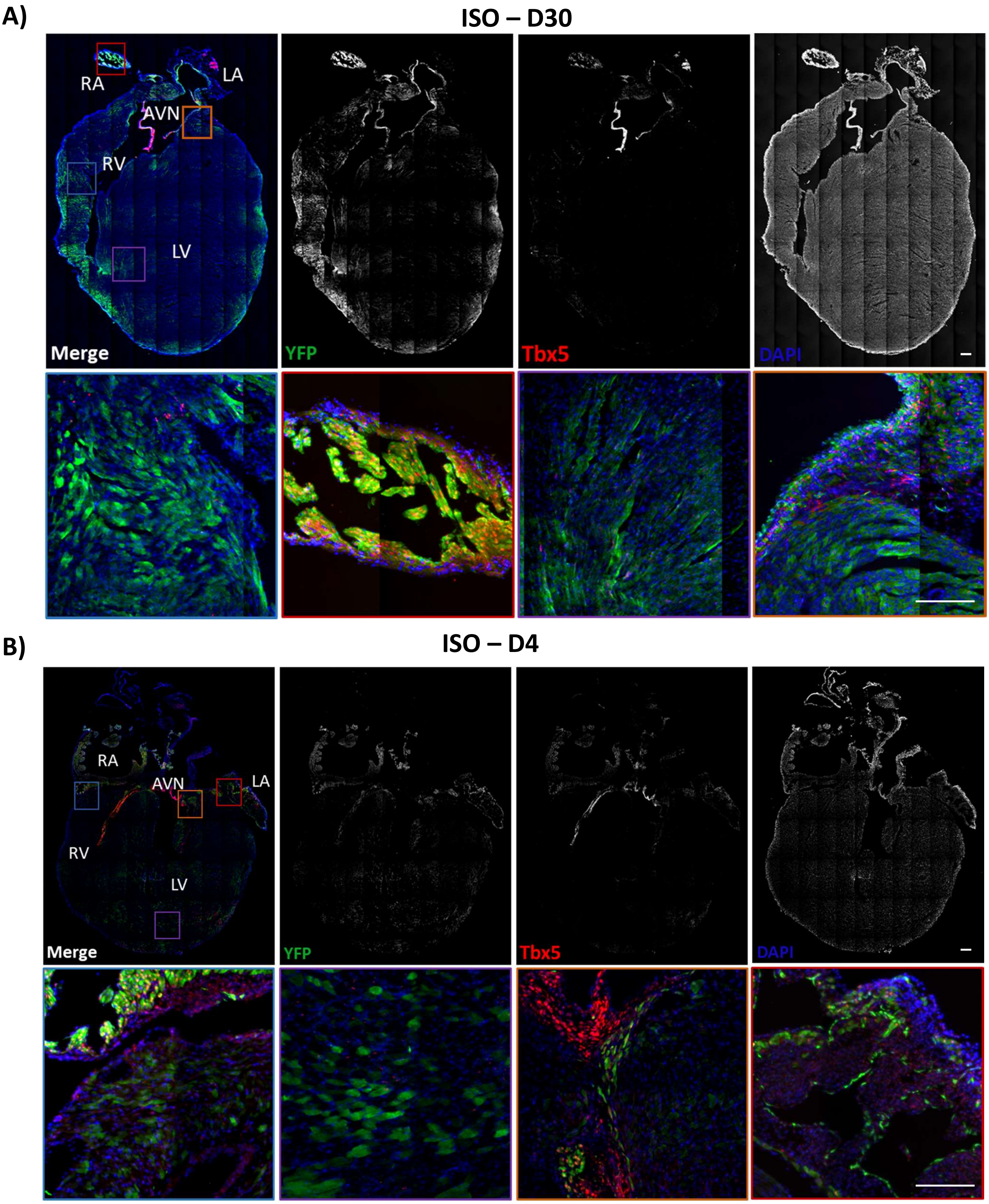


Supplementary Figure 4. (**A**) A collage of an ISO-injured adult heart on D30 after injury. (**B**) A collage of an ISO-injured adult heart on D4 after injury. Inserts indicating YFP$^{+}$Tbx5$^{+}$ are located in the atria on both D4 and D30, while YFP$^{+}$Tbx5$^{+}$ can be observed in D4 but not in D30 LV. N=1-3 hearts per timepoint. Key- LV=Left Ventricle, RV=Right Ventricle, RA=Right Atrium, LA=Left Atrium, AVN=Atrioventricular Node, SAN=Sinoatrial Node.

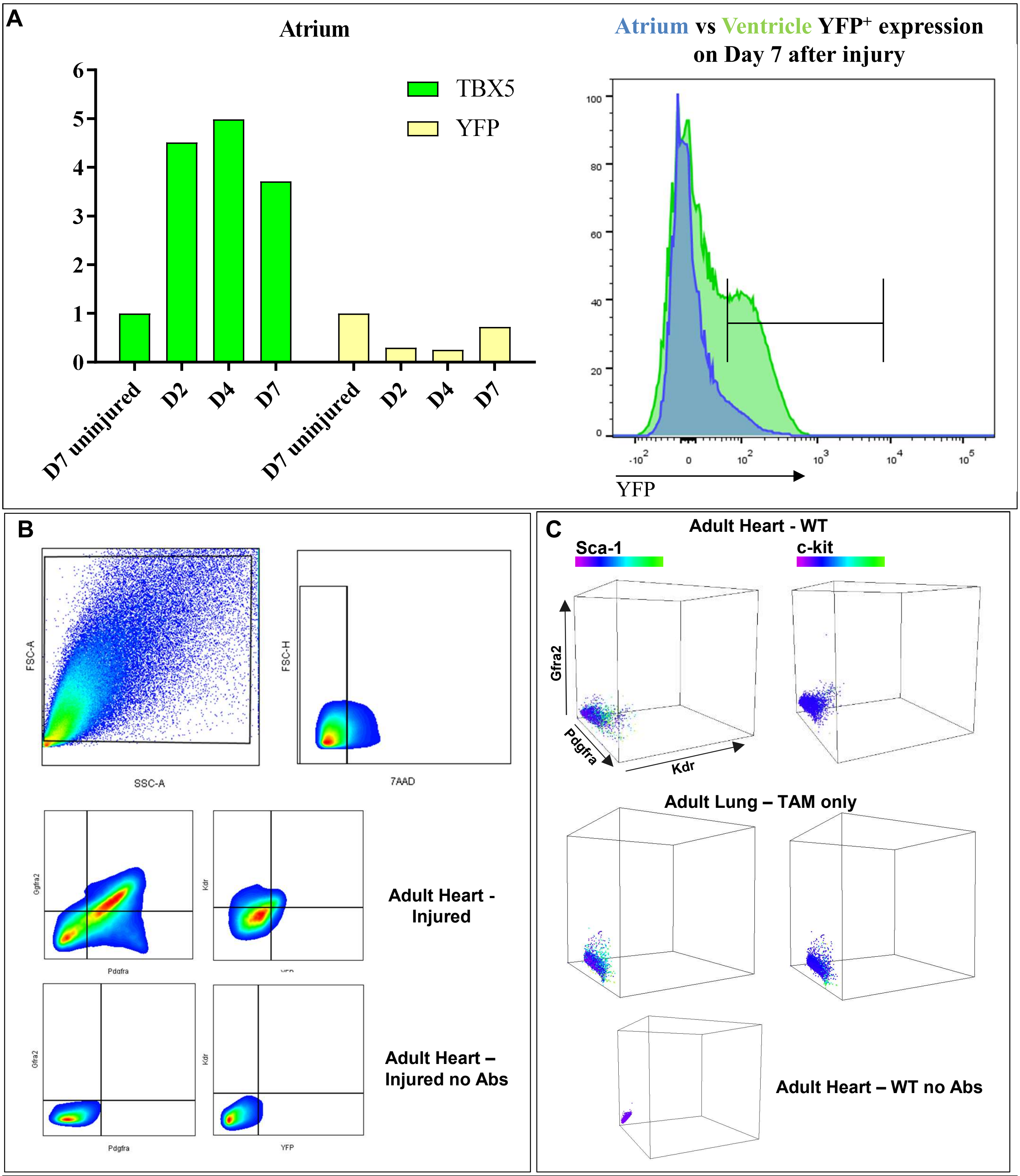


Supplementary Figure 5. (**A**) Real-time PCR analysis of *Yfp* and *Tbx5* transcripts in the atria of the adult heart in different time-points. Flow cytometry acquisition of YFP$^{+}$ cells from atria and ventricles seven days post-injury. (**B**) Representative graphs of flow cytometric gating strategy analysis from cardiac cells collected from adult injured hearts. (**C**) Representative graphs of flow cytometry acquisition of cardiac cells from uninjured heart and lung tissue.

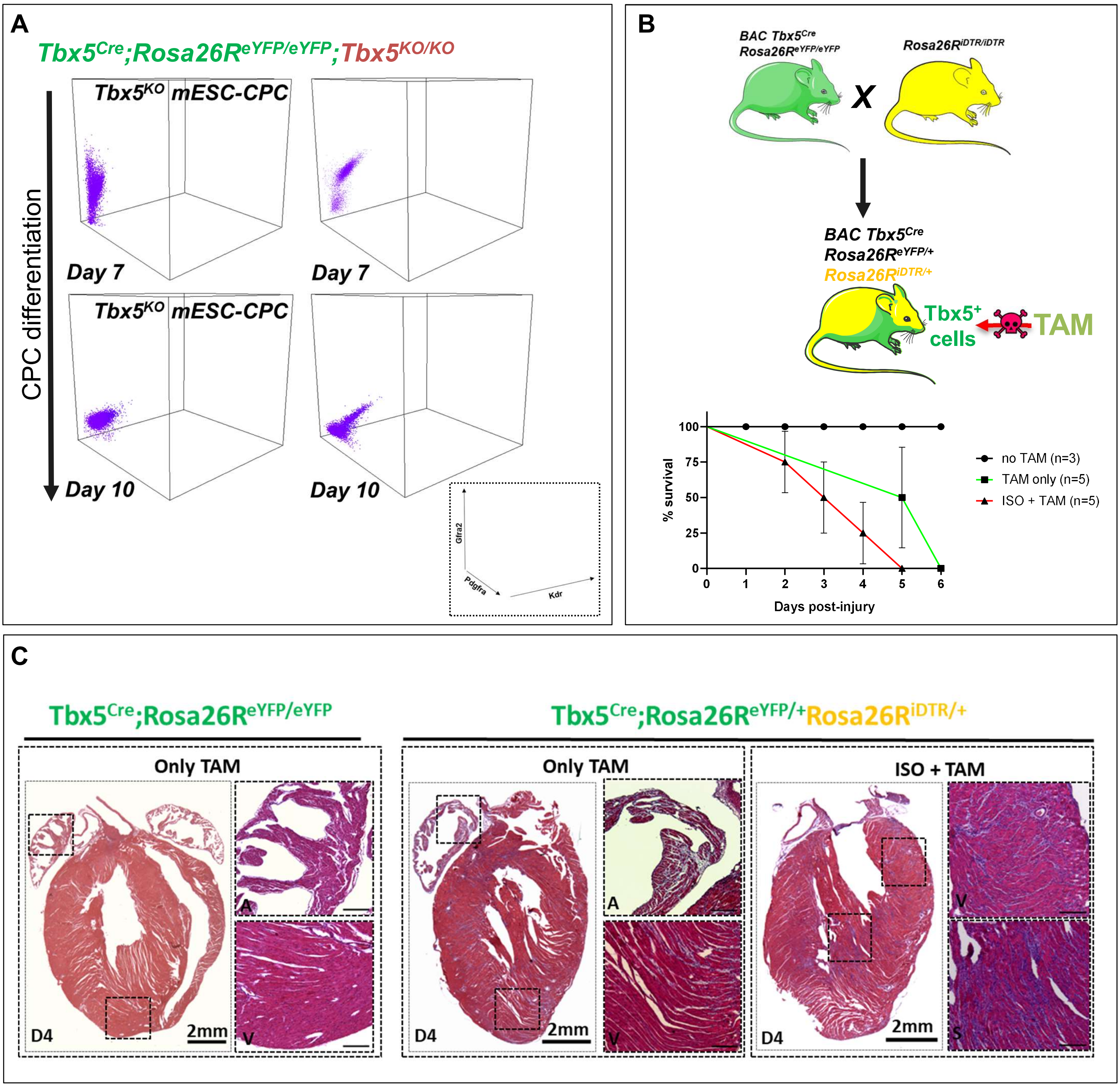


Supplementary Figure 6. (**A**) Representative three-dimensional graphs of flow cytometric acquisition of *Tbx5*$^{Cre}$*Rosa26R*$^{eYFP/+}$*Tbx5*$^{KO/KO}$-mESC CPC after 7 and 10 days under *in vitro* differentiating conditions. (**B**) *Tbx5*$^{Cre}$*Rosa26R*$^{eYFP/+}$*Rosa26R*$^{iDTR/+}$ cross and survival. Tamoxifen administration induces cells death to Tbx5$^{+}$-expressing cells. (**C**) Photos of adult hearts and haematoxylin and eosin staining confirmed non-fatal, yet substantial, heart injury in the left ventricle and atria, when compared to injured adult hearts. N=3 hearts per condition.

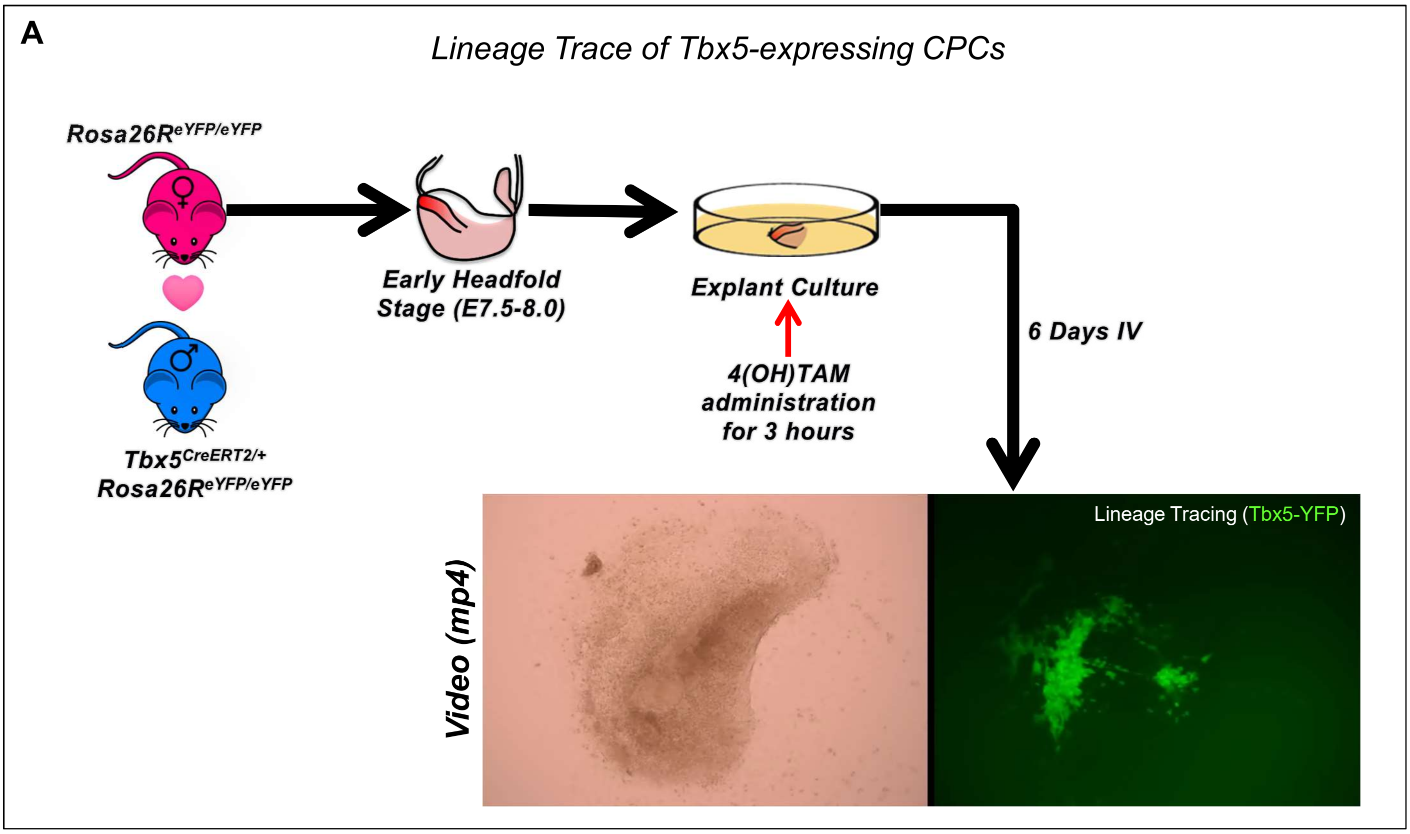


Supplementary Figure 7. Non-beating cardiac tissue was extracted from EHF stage embryos, show beating *ex vivo* in the areas where Tbx5 (YFP) was expressed. Video indicates the beating of $YFP^+$ cardiac cells.

**Supplementary Figure 8 – single-cell RNA-seq analysis QC** **Siatra *et al.***

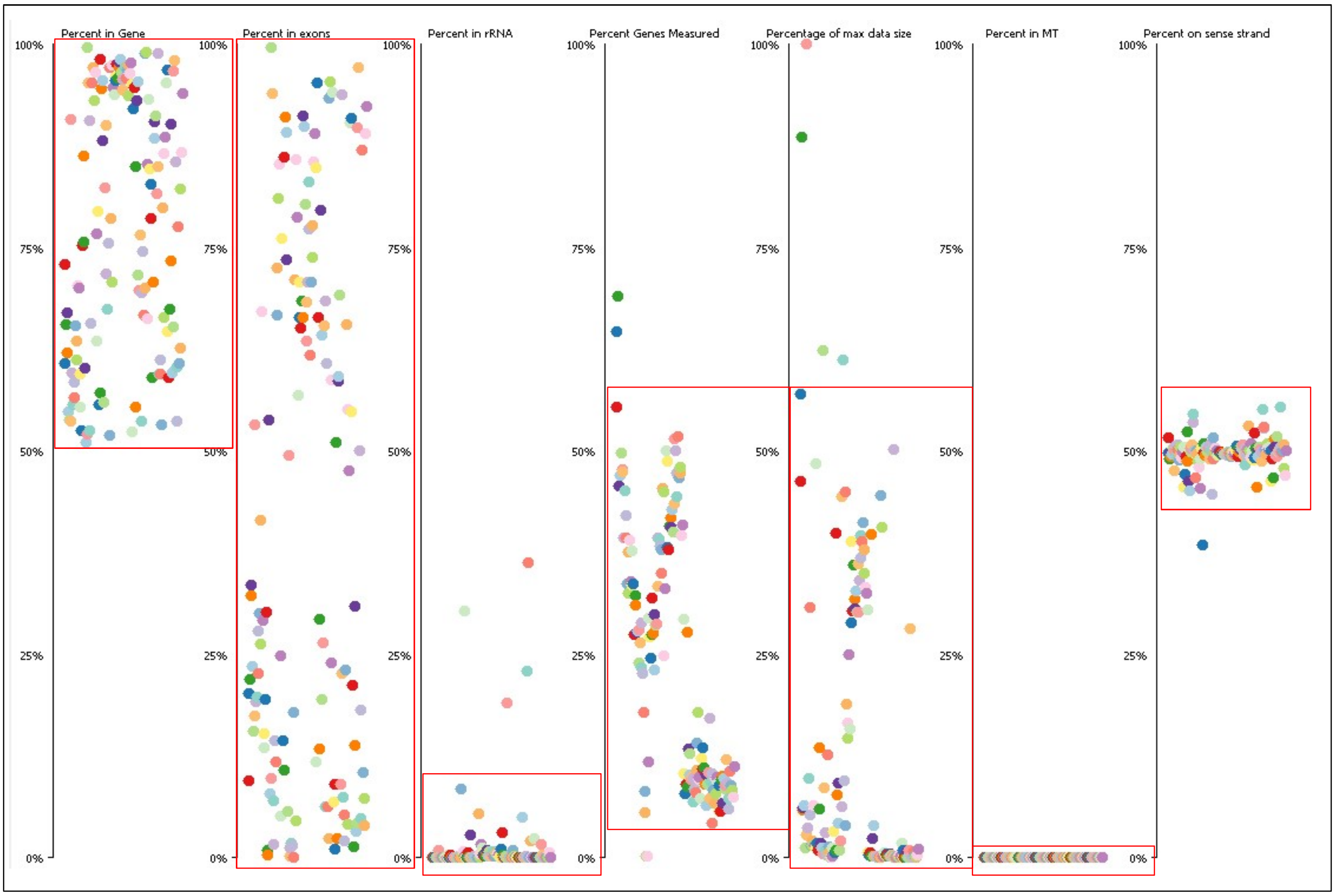


Supplementary Figure 8. Representative single-Cell RNA-seq QC report. In red, cut-off exclusion points for single-cells RNA-seq that have been chosen for downstream analysis (T-SNE, heatmaps, DEGs and GO/KEGG.

### Supplementary Table 1 Siatra *et al.*

| Input Parameter | Value |
|---|---|
| Single-end or paired-end reads | paired |
| Custom or built-in reference genome | indexed |
| Reference genome with or without an annotation | without-gtf |
| Select reference genome | mm10full |
| Gene model (gff3,gtf) file for splice junctions | |
| Length of the genomic sequence around annotated junctions | 100 |
| Use 2-pass mapping for more sensitive novel splice junction discovery | None |
| twopass_read_subset | Empty. |
| sj_precalculated | Empty. |
| Per gene/transcript output | - |
| Report chimeric alignments? | Don't report chimeric alignments |
| oformat | |
| Read alignment tags to include in the BAM output | NH (number of reported alignments/hits for the read) HI (query hit index) AS (local alignment score) nM (number of mismatches per (paired) alignment) ch (used to indicate chimeric alignments) |
| HI tag values should be | one-based |
| outSAMprimaryFlag | OneBestScore |
| MAPQ value for unique mappers | 60 |
| filter | |
| Exclude the following records from the BAM output | Nothing selected. |
| Would you like to set additional output filters? | yes |
| Would you like to keep only reads that contain junctions that passed filtering? | FALSE |
| Score range below the maximum score for multimapping alignments | 1 |
| Maximum number of alignments to output a read's alignment results, plus 1 | 10 |
| Maximum number of mismatches to output an alignment, plus 1 | 10 |
| Maximum ratio of mismatches to mapped length | 0.3 |
| Maximum ratio of mismatches to read length | 1 |
| Minimum alignment score | 0 |
| Minimum alignment score, normalized to read length | 0 |
| Minimum number of matched bases | 20 |
| Minimum number of matched bases, normalized to read length | 0 |
| Maximum number of multimapping alignments to output for a read | -1 |
| Calculation method for TLEN | leftmost base of the (+)strand mate to rightmost base of the (-)mate. (+)sign for the (+)strand mate |
| Configure seed, alignment and limits options | full |
| seed | |
| Search start point through the read | 30 |
| Search start point through the read, normalized to read length | 1 |
| Maximum length of seeds | 0 |
| Maximum number of mappings to use a piece in stitching | 10000 |
| Maximum number of seeds per read | 1000 |
| Maximum number of seeds per window | 50 |
| Maximum number of one seed loci per window | 10 |
| align | |
| Minimum intron size | 21 |
| Maximum intron size | 0 |
| Maximum gap between two mates | 0 |
| Minimum overhang for spliced alignments | 5 |
| Minimum overhang for annotated spliced alignments | 3 |
| Minimum mapped length for a read mate that is spliced | 0 |
| Minimum mapped length for a read mate that is spliced, normalized to mate length | 0.66 |
| Maximum number of windows per read | 10000 |
| Maximum number of transcripts per window | 100 |
| Maximum number of different alignments per read to consider | 10000 |
| Use end-to-end read alignments, with no soft-clipping? | FALSE |
| minimum number of overlap bases to trigger mates merging and realignment | 0 |
| maximum proportion of mismatched bases in the overlap area | 0.01 |
| chim_settings | |
| Minimum length of chimeric segment | 12 |
| Minimum total (summed) score of chimeric segments | 0 |
| Maximum difference of chimeric score from read length | 20 |
| Minimum difference between the best chimeric score and the next one | 10 |
| Penalty for a non-GT/AG chimeric junction | -1 |
| Minimum overhang for a chimeric junction | 20 |
| Maximum gap in the read sequence between chimeric segments | 0 |
| Discard chimeric alignments with Ns in the genome sequence around the chimeric junction | TRUE |
| Maximum number of multi-alignments for the main chimeric segment. | 10 |
| Maximum number of chimeric multi-alignments | 1 |
| Score range for multi-mapping chimeras | 1 |
| limits | |
| Maximum number of junctions for one read (including all multimappers) | 1000 |
| Maximum number of collapsed junctions | 1000000 |
| Maximum number of inserts to be inserted into the genome on the fly. | 1000000 |
| Number of genome bins for coordinate-sorting | 50 |

Supplementary Table 1. Parameters for aligning FASTQ raw data on the mm10 mouse genome using the RNA-STAR tool in http://www.usegalaxy.eu
